# Preclinical Study of Cannflavins A and B Action Against Glioblastoma Cells

**DOI:** 10.1101/2025.07.02.662823

**Authors:** Jennifer Holborn, Hannah Robeson, Ellis Chartley, Tiana Gluscevic, Adina Borenstein, Colby Perrin, Begüm Alural, Himain Perera, Tariq A. Akhtar, Nina Jones, Jasmin Lalonde

## Abstract

**Background:** Flavonoids represent a large group of naturally occurring polyphenolic compounds, many of which have been found to produce valuable biological outcomes, including action against cancer cells. Glioblastoma multiforme (GBM) is an aggressive type of brain tumor that is associated with a poor prognosis and has limited treatment options. Previous findings suggest that cannflavins can produce promising effects for pancreatic and bladder cancers, but the efficacy of these phytochemicals in attacking brain tumour cells remains unknown. Our study evaluates the potential of cannflavin A and cannflavin B against GBM cells using a range of *in vitro* approaches.

**Results:** We conducted experiments using A-172 and U-87 GBM lines to assess the impact of cannflavins A and B on cell viability, cycle, migration, and invasion capacity. Our results revealed a consistent dose-dependent decrease in cell viability in both lines after the addition of cannflavin B to the culture media. Interestingly, we found that chrysoeriol (the non-prenylated synthesis precursor of cannflavins in the *Cannabis sativa* plant) only has a limited adverse effect on survival using the same approach, while techtochrysin (an O-methylated flavone) has none. Using time-lapse live-cell imaging and a scratch assay, we also show that cannflavins can inhibit tumor cell migration at concentrations below those that produce significant cell death. Finally, we found that cannflavin B exhibits anti-migratory and anti-invasive properties in transwell and tumorsphere assays, underscoring its multifaceted therapeutic potential.

**Conclusion:** These findings suggest that cannflavin B holds promise as a therapeutic agent in the treatment of GBM.

**Highlights:**

- Cannflavin B but not cannflavin A limits the viability of A-172 and U-87 GBM cells in a concentration- and time-dependent manner.
- Using time-lapse live-cell imaging and a scratch assay, we provide evidence that cannflavins A and B can inhibit GBM cell migration at concentrations below those that result in a significant reduction in cell viability.
- Cannflavin B exhibits robust anti-migratory and anti-invasive properties in transwell and tumorsphere assays, underscoring its multifaceted therapeutic potential.
- Although previous work had reported the capacity of cannflavins A and B to interfere with TrkB and downstream the MAPK/AKT signaling pathways in mouse primary neurons, the effect of those molecules is limited in GBMs, suggesting that other targets are engaged to produce the cellular effects.

## Introduction

Flavonoids are naturally occurring polyphenolic molecules that accumulate in dietary items such as fruits, vegetables, and teas. This broad group of phytochemicals has received considerable research attention for over 25 years due to their beneficial anti-inflammatory and anti-oxidative properties (Panche *et al*., 2016). Many flavonoids have also attracted interest in cancer research due to their potential therapeutic applications. For instance, kaempferol has been found to induce G2/M cell cycle arrest as well as reduce migration and invasion in breast, liver, ovarian, lung, and brain cancers (Gao et al., 2018; Jo et al., 2015; Li et al., 2017; Zhu et al., 2019). Apigenin and luteolin, for their part, have shown promise as anti-cancer agents for breast, prostate, lung, and brain subtypes by suppressing the mitogen-activated protein kinase (MAPK) and Protein kinase B (AKT) signaling pathways that control cell proliferation and survival (Cheng et al., 2013; Erdogan et al., 2016; Huang et al., 2019; Tseng et al., 2017). Finally, quercetin, one of the most consumed dietary flavonoids, induces G2/M phase arrest and significantly increases tumor suppressor protein p53 expression in MDA-MB-453 breast cancer cells (Choi et al., 2008). It also limits angiogenesis and metastasis of other cancer cell types through changes in MAPK/AKT/mTOR signaling pathways (Balakrishnan et al., 2016; Gao et al., 2012; Kim et al., 2019; Lee et al., 2015; Pratheeshkumar et al., 2012). Taken together, these examples support searching for additional flavonoids that also produce robust anti-cancer effects.

Cannflavins are prenylated and highly lipophilic metabolites first purified 40 years ago from *Cannabis sativa (C. sativa)* plant extracts (Barrett et al., 1985). As illustrated in **Supplementary Figure 1**, the cannflavin family includes five known members, which are cannflavin A, cannflavin B, cannflavin C, isocannflavin B (also known as FBL-03G), and 8-prenylcannflavin B (also known as dorsmanin D). Several of these molecules have been shown to produce promising effects applicable to various clinical conditions. For instance, studies revealed the anti-inflammatory properties of cannflavins A and B through their ability to inhibit pro-inflammatory mediators and the production of prostaglandin E_2_ at a rate thirty times more potent than aspirin (Barrett et al., 1985; Werz et al., 2014). Consistent with their anti-inflammatory activity, Eggers and colleagues (2019) have reported neuroprotective action of cannflavin A in neuronal PC12 cells against amyloid β-induced cell death by reducing Aβ_1_-42 aggregation and fibril formation.

Beyond studies concerning inflammation, other research suggests that specific cannflavin molecules exhibit desirable anti-cancer effects that may involve other pathways. For instance, isocannflavin B was found to improve survival in preclinical mouse models of metastatic pancreatic cancer by increasing apoptosis of tumor cells (Moreau et al., 2019), and cannflavin A has been reported to synergize with agents commonly used to treat bladder cancer at specific concentrations (Tomko et al., 2022). Despite those promising results, whether cannflavins can effectively attack brain cancer cells has not been directly studied.

Glioblastoma multiforme (GBM) is the most aggressive and common primary brain tumor in adults, characterized by rapid growth and diffuse infiltration into surrounding healthy brain tissues (Tilak et al., 2021a). The prognosis for GBM patients is extremely poor, with a median survival of approximately 15 months despite aggressive treatment (Johnson and O’Neill, 2012; Ostrom et al., 2020). The standard of care for GBM is maximum safe surgical resection followed by a combination of radiation therapy and temozolomide (TMZ) chemotherapy, a DNA alkylating agent that induces cell death through DNA lesions (Stupp et al., 2005; Stupp et al., 2009; Fisher and Adamson, 2021). However, the inability to fully resect the tumor with surgery, combined with the persistence of TMZ-resistant cancer cells capable of recapitulating tumors, typically leads to disease relapse within just a few months (Weller et al., 2013; Birzu et al., 2020). Therefore, there is a pressing need to identify new agents that could improve patient prognosis.

In this study, we conducted experiments using the A-172 and U-87 GBM cell lines to assess the effects of cannflavins A and B on cell viability, migration, and invasion. Our work revealed a decrease in GBM cell viability with increasing concentrations of cannflavin B. Additionally, analysis of cell migration using a time-lapse live-cell scratch assay revealed the potential of cannflavins to impede tumor dissemination at concentrations below those inducing cell death. Specifically, cannflavin B exhibited robust anti-migratory and anti-invasive properties, as evidenced by transwell and tumorsphere invasion assays, highlighting its multifaceted therapeutic effects. Finally, as our previous work found interference of TrkB signaling by cannflavin A and cannflavin B in mouse primary neurons (Holborn et al., 2023), we also evaluated whether the action of cannflavins on GBM cells could be tied to reduced activity of this receptor tyrosine kinase (RTK). Together, our findings underscore the promising anti-cancer properties of cannflavin B against immortalized GBM cells, suggesting its potential role as a novel therapeutic agent.

## Materials and methods

### Cell culture and reagents

Malignant male glioma cell lines A-172 and U-87 MG (U-87) were obtained from the American Type Culture Collection (ATCC, USA, Cat No. CRL-1620 and HTB-14, respectively). Cells were cultured in DMEM (Gibco, USA, Cat No. 11965-092) supplemented with 10% FBS (Gibco, Cat No. 12483-020) and 0.5% penicillin/streptomycin (Gibco, Cat No. 155140-122), at 37°C and 5% CO_2_. For treatment with cannflavins and other tested agents, the molecules were dissolved in DMSO and applied directly to the culture media.

The BIONi010-A cell line (European Bank for induced Pluripotent Stem Cells, ECACC Cat No. 66540022) used to prepare cerebral organoids was purchased from Sigma-Aldrich (St-Louis, MO, USA) and cultured in ExCellerate iPSC Expansion Medium (R&D Systems, Cat No. CCM036) according to the manufacturer’s instructions.

### Antibodies and pharmacological compounds

The antibodies recognizing phosphorylated p44/p42 MAPK (Cat No. 4370), AKT (Cat No. 4691), phosphorylated AKT Thr 308 (AKT T308, Cat No. 2965), and phosphorylated AKT Ser 473 (AKT S473, Cat No. 4060) were acquired from Cell Signaling Technology (Beverly, MA, USA). The antibody detecting phosphorylated TrkB Tyr705 (TrkB Y705, Cat No. ab229908) was from Abcam (Waltham, MA, USA), the antibody recognizing p42 MAPK (Erk2, sc-1647) was from Santa Cruz Biotechnology (Santa Cruz, CA), the GAPDH (Cat No. AB2302) antibody was from Sigma-Aldrich, and the Tubulin β 3 (TUJ1, Cat No. 801202) antibody was from BioLegend (San Diego, CA, USA). Finally, the GFP (Cat No. 10262) and cross-absorbed secondary antibodies (horseradish peroxidase- and fluorophore-conjugated) were from Invitrogen.

Cannflavin A (Cat No. C-291001) and cannflavin B (Cat No. C-291003) were purchased from TLC Pharmaceutical Standards (Newmarket, ON, Canada). ANA-12 (Cat No. SML0209) was acquired from Sigma-Aldrich, and chrysoeriol (Cat No. 491-71-4) from Cayman Chemical (Ann Arbor, MI, USA).

Techtochrysin was biosynthesized as follows: The *O. basilicum* O-methyl transferase ObFOMT1 (Accession number JQ653275) open reading frame was synthesized by Twist Bioscience (San Francisco, CA, USA) in the *pET28a* vector system between the *NdeL* and *XhoI* sites. This system inserted an N-terminal hexahistidine tag into the coding sequence. The construct was then introduced into *E. coli* BL21-CodonPlus (DE3)-RIPL. Bacterial cells harboring the expression vector were cultured in LB media at 37°C to an OD_600_ of 0.6. Isopropyl-β-D-thiogalactoside was added to a final concentration of 0.2 mM, and cultures were further incubated at 16°C for an additional 16 hours to allow for protein expression. Thereafter, *E. coli* cells were collected by centrifugation, resuspended in Buffer A (20 mM Tris-HCl, 500 mM KCl, pH 8.0) and lysed using an Emulsiflex. Cell debris and remaining intact cells were removed from the solution by centrifugation (20,000 x *g*, 20 minutes at 4°C) and the supernatant was applied to a 1 mL HisTrap FF column (Cytiva, Marlborough, MA) equilibrated in Buffer B (20 mM Tris-HCl, 500 mM KCl, 20 mM imidazole, pH 8.0). Proteins bound to the Ni^2+^ affinity matrix were washed with fifteen column volumes of Buffer B, eluted with one column volume of Buffer C (20 mM Tris-HCl, 500 mM KCl, 200 mM imidazole, pH 8.0), and then immediately desalted on PD-10 columns (Cytiva) equilibrated with Buffer D (50 mM Tris-HCl, 5 mM MgCl_2_, pH 8.0, 10% glycerol). The concentration of recombinant ObFOMT1 was determined by Bradford Assay using BSA as a standard, and 1 mg was used per enzymatic assay to produce techtochrysin. Precisely, purified recombinant ObFOMT1 was mixed in 10 mL of assay solution (1 mM chrysin, 10% w/v hydroxypropyl-β-cyclodextrin, 2 mM S-adenosylmethionine, and 5 mM MgCl_2_ in 50 mM Tris-HCl, pH 8.0) and incubated with shaking for 16 hours at 15°C. Reagents for the assay solution were obtained from Xi’an B-Thriving I/E Co., Ltd (Xi’an, Shaanxi, China). The enzymatic reaction was terminated by adding 1 mL of 10% formic acid. Prenylated bibenzyl products were extracted with three volumes of ethyl acetate three times, evaporated to dryness with gaseous N, and resuspended in 6 mL of methanol. The sample was chromatographed on an Infinity Lab Poroshell 120 EC-C18 2.7 μM (ODS2 9.4 x 150 mm) column and resolved by an Agilent 1260 infinity HPLC using a linear gradient with solvent A (45% methanol, 0.1% formic acid in water) and solvent B (95% methanol, 0.1% formic acid in water) at a flow rate of 2.5 mL/min heated at 30°C. The gradient started at 60% of solvent B and increased to 70% in 7 minutes, then increased to 100% of solvent B in 1 minute and remained isocratic for 2 minutes. Techtochrysin was detected by absorption at 340 nm and was collected from the HPLC at 9.3 minutes. Finally, the eluate was dried under gaseous nitrogen, lyophilized, and techtochrysin quantified relative to a standard curve generated with chrysin. Product identity was confirmed based on mass spectral fragmentation patterns.

### Cell viability assay

Cell viability was assessed using a resazurin reduction assay, as described previously (Holborn et al., 2023). Briefly, A-172 and U-87 cells were seeded in 96 multi-well plates at a density of 1.0 × 10^4^ per well and allowed to attach overnight. Cells were then treated with DMSO and varying concentrations of ANA-12 (1 to 50 µM) or cannflavins (1 to 20 µM) diluted in Opti-MEM reduced serum media (Gibco, Cat No. 31985070) and incubated at 37°C for the indicated time. Following treatment, resazurin sodium salt (Sigma-Aldrich, Cat No. R7017) was dissolved in Hank’s balanced salt solution (Wisent Inc., Saint-Jean-Baptiste, QC, Canada, Cat No. 311-513-CL), diluted 1:10 in each well, and cells were further incubated at 37°C for three hours. Fluorescence was measured on a microplate reader at 570/590 nm, and a background control was subtracted from each value. All treatments were completed in triplicate.

The following procedure was used for quantification: First, fluorescence values were exported from the microplate reader, and the average of three background control wells with Opti-MEM reduced serum media only was subtracted from each treatment value. Second, the subtracted values were multiplied by 100 and divided by the average of the DMSO controls to obtain a percentage relative to the vehicle control. Finally, the average of three replicates per treatment was used as the final value for one biological replicate. A minimum of three biological replicates were taken.

### Lactate dehydrogenase cytotoxicity assay

Lactate dehydrogenase (LDH) release was measured using the CyQUANT LDH Cytotoxicity Assay Kit according to the manufacturer’s protocol (Invitrogen, Cat No. C20300). Briefly, A-172 and U-87 cells were seeded in 96-well multiplates and treated with the tested molecules, using a similar approach to the resazurin reduction assay. Before the treatment endpoint, 10× Lysis Buffer was applied to control wells for 10 minutes to get the Maximum LDH Release value. Cell culture supernatant from each treatment was transferred to a new 96-well multiplate, an equal volume of reaction mixture was added, mixed, and then the plate was incubated at room temperature for 30 minutes, protected from light. Absorbance was measured on a microplate reader at 490 nm and 680 nm (background). All treatments were completed in triplicate.

The following procedure was used for quantification: First, LDH activity was calculated by subtracting the raw absorbance value measured at 680 nm from the 490 nm value for each replicate and treatment. The average of three background control wells with Opti-MEM reduced serum media was subtracted from each treatment value. Finally, the LDH activity of each technical replicate of the compound-treated wells was divided by the average of the maximum LDH activity and multiplied by 100 to obtain the percent cytotoxicity. The average of the triplicates per treatment was taken and used as the final value for one biological replicate; three biological replicates were completed in total.

### Terminal deoxynucleotidyl transferase dUTP nick end labeling (TUNEL) assay

The Roche *in Situ* Cell Death Detection Kit, Fluorescein (Sigma-Aldrich, Cat No. 11684795910), was used according to the manufacturer’s protocol. Briefly, A-172 cells were fixed using 4% paraformaldehyde for 15 minutes after treatment, then incubated for 2 minutes in permeabilization solution (0.1% Triton X-100, 0.1% sodium citrate in PBS). This was followed by a 1-hour incubation in the dark at 37°C with humidity in TUNEL reaction solution. After rinsing with PBS three times for 10 minutes, cells were incubated in 4’,6-diamidino-2-phenylindole (DAPI; Invitrogen, Cat No. 62247) staining solution for 10 minutes and finally washed in PBS twice for 10 minutes before being mounted onto slides with ProLong Diamond Antifade (Invitrogen, Cat No. P36930).

### Flow cytometry

A-172 cells and U-87 cells were seeded in 6-well plates at a density of 2.5 × 10^5^ per well and allowed to attach overnight. The next day, cells were treated with DMSO control, cannflavin A, or cannflavin B (1 and 5 µM) for 24 hours. Following that time, the cell monolayer was dissociated with trypsin, pelleted, resuspended in FACS buffer (PBS supplemented with 2% FBS and 1 mM EDTA), and frozen fixed at −20°C in 70% ethanol. Immediately before running samples, fixed cells were thawed and washed twice in FACS buffer, 50 µL of 100 µg/mL RNAse A (Thermo Fisher Scientific, Cat No. EN0531) in FACS buffer was added for 30 minutes, then 200 µL of 0.25 µg/mL propidium iodide (Molecular Probes, Cat No. P-1304) in FACS buffer was added for 10 minutes in the dark. The DNA contents were analyzed using a Northern Lights cytometer (Cytek Biosciences, Bethesda, MD, USA).

Data quantification was performed using FlowJo v10.10 software. First, debris was excluded using SSC-A vs. FSC-A gating strategy, and then singlets were selected using FSC-H vs. FSC-A. Specifically, the univariate cell cycle platform Watson Pragmatic algorithm distributed cell cycle peaks to the histograms presented, and data points were plotted as a percentage of the cell count found in each cell cycle phase (G1, S, G2/M).

### Transwell invasion assay

The bottoms of Corning Transwell inserts (New York, USA, Cat No. CLS3442) were coated with 20 µg/mL fibronectin (Sigma-Aldrich, Cat No. F1141-1MG) at 4°C for 1 hour. After three washes with PBS, the tops were coated with 0.2 mg/mL Corning Matrigel matrix (Cat No. 354234) overnight at 37°C. In parallel with the preparation of the transwells, A-172 cells were seeded at a density of 1.5 × 10^5^ cells per well in 6-well plates and grown to 85-90% confluency for 24 hours. They were then treated overnight with 10 µM cannflavin B or DMSO in DMEM containing 1% FBS media. The next day, 30,000 live cells from each condition were collected and seeded in the upper chamber of each well with DMEM containing 1% FBS plus cannflavin B (10 µM), and in the bottom chamber with DMEM containing 10% FBS. Cells were allowed to invade the Matrigel matrix for 24 hours at 37°C and 5% CO_2_. Afterward, the transwell membranes containing the invaded cells were removed. The membranes were stained with a 1% crystal violet solution in 20% methanol for 20 minutes, washed three times with PBS, and dried for several hours at room temperature. Representative photographic images of the membranes were taken and then cut out. Finally, crystal violet was eluted from each membrane with 10% acetic acid for 20 minutes, and the absorbance (at 562 nm) of the eluates was measured to quantify invasion. Three biological replicates with each treatment run in duplicate were analyzed.

### Scratch migration assay

A-172 cells were seeded in a 6-well plate at a density of 2.0 × 10^5^ per well and grown to complete confluency in DMEM supplemented with 10% FBS for 48 hours. Using a P1000 pipette tip, 4 crossing linear wounds were created through the monolayer of cells within each well. Next, the unattached cells were washed away with PBS twice before being added back to 1% FBS DMEM supplemented with DMSO only or the treatment molecules. Baseline images of the wound area were acquired immediately after adding the culture media using a phase-contrast inverted Nikon Eclipse Ti2-E microscope (Nikon Instruments, Melville, NY, USA) equipped with a motorized stage and STXF stage top incubator (Tokai Hit USA Inc., PA, USA) to maintain temperature (37°C), humidity, and CO_2_ (5%, 150 mL/min) constant. Two separate areas per scratch were imaged every 15 minutes over a 48-hour period. NIS-Elements Advanced Research software was used to run binary area fraction analysis, and the number of migrated cells was counted using ImageJ v1.54 software. Three biological replicates were performed for each treatment condition.

### Generation of tumorspheres (tumor spheroids)

The tumorsphere protocol was adapted from previous work (Vinci et al., 2015; Tilak et al., 2021b). Briefly, U-87 cells were passaged at least three times in complete media and then 20,000 cells per well were seeded into a clear round-bottom ultra-low attachment microplate (Corning, Cat No. 7007). Tumorspheres were allowed to form for 5 days at 37°C and 5% CO_2_, with half the media changed on Day 3.

### Tumorsphere invasion assay

Individual tumorspheres prepared as mentioned above were embedded in Corning Matrigel matrix (Cat No. 354234) containing the specified treatment. Four tumorspheres with a circular shape and no protrusions were chosen per treatment, and half the media was removed from the well. An equal volume of Matrigel supplemented with treatment at 2x concentration was added to each well and incubated at 37°C and 5% CO_2_ for one hour to polymerize. Treatments at 1x of the desired treatment concentration were prepared in complete media and added to each well immediately after incubation. The plate was then imaged on a Nikon Eclipse Ti2-E inverted microscope equipped with a motorized stage, image stitching capability, and a 10× objective at 0, 24, 48, and 72 hours. One image per tumorsphere per treatment (4 tumorspheres and 4 treatments for 16 images at each time point) was analyzed, and three biological replicates were performed.

The image analysis on original tiff files was performed with ImageJ as follows: For the area measurement, the elliptical tool was used to encircle each tumorsphere, ensuring all cells grown out from the original tumorsphere were included in the area measurement. Three area measurements were taken for each tumorsphere at each time point and averaged. For the radius measurement, the line tool was used to measure from the center point of the tumorsphere out to the furthest protrusion from the tumorsphere. Three radius measurements were taken for each tumorsphere at each time point and averaged. The average area or radius of each treatment was divided by the DMSO control average at its corresponding time point to obtain the values used for statistical analysis.

### Generation of cerebral organoids

BIONi010-A induced pluripotent stem cells (iPSCs) were cultured on Matrigel-coated plates (Corning, Cat No. 354277) in ExCellerate iPSC Expansion Medium (R&D Systems, Cat No. CCM036). Cerebral organoids were generated using the STEMdiff Cerebral Organoid Kit (StemCell Technologies, Cat No. 08570) as described in the manufacturer’s protocol and Tilak and colleagues (2021b). Briefly, on Day 0, iPSCs were lifted with Gentle Cell Dissociation Reagent, resuspended in embryoid body (EB) Seeding Medium, and seeded in an ultra-low attachment, round bottom 96-well plate (Corning, Cat No. 7007) at a density of 9,000 cells/well. EB Formation Medium was added on Days 2 and 4, and on Day 5, EBs were transferred to ultra-low attachment 24-well plates (Corning, Cat No. 3473) containing Induction Medium for 48 hours. On Day 7, EBs were encased in droplets of Matrigel on organoid embedding sheets (StemCell Technologies, Cat No. 08579), which were allowed to polymerize at 37°C before being washed into ultra-low attachment 6-well plates (Corning, Cat No. 3471) using 3 mL of Expansion Medium. On Day 10, the Expansion Medium was replaced with Maturation Medium, and the plate was transferred to an orbital shaker inside a 37°C, 5% CO_2_ incubator. The media was changed three times per week until Day 60, the beginning of the organoid co-culture assay.

### Organoid co-culture assay

The organoid co-culture assay protocol was modified from Tilak and colleagues (2021b). Tumorspheres were generated similar to above, except with 12,000 GFP-expressing U-87 cells seeded per well. Sixty (60)-day-old organoids were removed from the 6-well plate and placed individually in 1.5 mL Eppendorf tubes, each containing 4 tumorspheres and 500 μL of Maturation Medium. Tubes were incubated at 37°C and 5% CO_2_ inside 10 cm dishes with the caps open to allow airflow during attachment, while maintaining a sterile environment. After 48 hours, 50% of the media was replaced, and after 72 hours, organoids with tumorsphere attached were gently removed from the tubes using a wide-bore pipette and put back in 6-well plates on the orbital shaker for an additional 24 hours (for the 10-day experiment) or 8 days (for the 3-day experiment), at which point cannflavin B (5 μM) or DMSO was added to the well in a complete media change. Media containing cannflavin B or DMSO was replaced every other day for the duration of the experiment. Fourteen days after initiating the co-culture, all organoids were removed from the wells, washed three times with PBS, and fixed in 4% paraformaldehyde overnight at 4°C. Organoids were washed again with PBS and incubated in a 30% sucrose solution at 4°C for 4 days before being embedded in a 7.5% gelatin, 10% sucrose solution and snap-frozen in dry ice submerged in 95% ethanol. Twenty (20) μm sections were obtained using a cryostat (Leica, CM1860), placed on Vectabond-coated slides (Vector Laboratories, Cat No. SP-1800-7), and stored at −80°C until immunostaining.

For immunostaining, organoid sections were washed 3 times with PBS and blocked in 10% normal goat serum in PBS at room temperature for 30 minutes. Sections were incubated overnight at 4°C in blocking solution containing primary antibodies (TUJ1 and GFP, 1:1,000). Slides were washed again 3 times with PBS and incubated in Alexa Fluor-488- or Alexa Fluor-594-conjugated secondary antibodies (Invitrogen, Cat Nos. A11039 and A11032) in 5% normal goat serum in PBS for 2 hours at room temperature. Slides were washed 3 additional times in PBS, counterstained with DAPI, and cover-slipped with ProLong Glass Antifade mounting reagent (Invitrogen, Cat No. P36984).

Captures were taken using a Nikon Eclipse Ti2-E microscope at 20× magnification. Exposure for each channel was kept consistent between conditions. To analyze GFP-positive U-87 cell infiltration in the corresponding cerebral organoid, the following steps were followed: 1) the perimeter of the cerebral organoid was traced based on the TUJ1 signal, 2) a threshold of 100 was applied to the GFP channel to binarize pixels, and 3) the number of GFP pixels was normalized to the total number of pixels in the TUJ1 channel to provide the proportional count. For estimating tumorsphere area, a section from approximately the middle of each tumorsphere to be analyzed was immunostained, and the perimeter of the GFP signal was traced using the polygon selection tool in ImageJ software by an individual blinded to the experimental conditions.

### Western blotting

For western blot analysis, cells were seeded in 6-well plates and grown for 2 to 3 days before treatment. After treatment, cells were collected with radioimmunoprecipitation assay (RIPA) buffer (50 mM Tris [pH 8.0], 0.5% Igepal-CA-630, 0.1% SDS, 0.5% deoxycholic acid, 1 mM EDTA, 300 mM NaCl), supplemented with Phosphatase Inhibitor Cocktail 3 (Sigma-Aldrich, Cat No. P0044) and EDTA-free Protease Inhibitor Cocktail (Sigma-Aldrich, Cat No. 11836170001), and homogenized using QIAshredder columns (QIAGEN, Cat No. 79656). Concentration of all lysates was measured using Pierce BCA Assay Kit (Thermo Fisher Scientific, Cat No. 23225), one volume of 2× Laemmli buffer (100 mM tris-HCl [pH 6.8], 4% SDS, 0.15% bromophenol blue, 20% glycerol, 200 mM β-mercaptoethanol) was added, samples were adjusted to an equal concentration, and finally boiled for 5 minutes. A total of 15 µg of protein was run on a 10% SDS-PAGE gel and transferred to a nitrocellulose membrane. Membranes were incubated in TBST (Tris-buffered saline and 0.1% Tween 20) supplemented with 5% non-fat powdered milk for 30 min at room temperature, and then probed overnight with the intended primary antibody at the following dilution: P-TrkB Y705 (1:1,000), P-AKT S473 (1:1,000), P-AKT T308 (1:1,000), pan-AKT (1:2,000), P-p44/p42 MAPK (1:1,000), p42 MAPK (1:1,000), and GAPDH (1:20,000). The next day, membranes were washed with TBST, incubated with the appropriate secondary antibody, and visualized using enhanced chemiluminescence (ECL), according to the manufacturer’s guidelines (Pierce, Thermo Fisher Scientific). Each experiment was repeated three times.

The western blots were quantified as follows: First, equal amount of protein sample was analyzed by SDS-PAGE for each biological replicate. Second, the exposure time of the film to ECL was the same for each biological replicate. Third, the exposed films were scanned using a HP LaserJet Pro M377dw scanner in grayscale at a resolution of 300 dpi. Finally, protein expression was quantified using ImageJ software to determine the mean grey value of the protein bands, with the background value subtracted. The pixel density for all data was inverted by subtracting the mean grey value from 255. The final values of each protein band were taken as a ratio over the corresponding loading control to provide a normalized value for statistical analysis.

### Statistical analysis

All statistical calculations were completed with Prism 10 (GraphPad Software, Boston, MA, USA). No statistical methods were employed to predetermine the sample size of any of the presented experiments. Student *t*-test or analysis of variance (ANOVA) plus Tukey’s post hoc test for multiple comparisons was performed where indicated. A value of *p* ≤ .05 was considered statistically significant. No test for outliers was conducted, and no data point was excluded. Unless mentioned otherwise, all results represent the mean ± SEM from at least three independent experiments (*n* = 3). Sample size was estimated based on previous studies of a similar nature (Lalonde et al., 2021; Holborn et al., 2023; Lau et al., 2025).

## Results

### Cannflavin B decreases GBM cell viability

The flavonol kaempferol and flavone apigenin have been shown to significantly reduce the viability of different cancer cell types in the low micromolar concentration range after 24-48 hours of exposure (e.g., Gao *et al*., 2018; Tseng *et al*., 2017). Except for the work of Moreau and colleagues (2019) using mouse models of metastatic pancreatic cancer and the study by Tomko and colleagues (2022), which reported a promising effect of cannflavin A against bladder carcinoma cells, we know little about the impact of cannflavins on other cancer types. This prompted us to thoroughly test the effect of cannflavin A and cannflavin B on the viability of the A-172 and U-87 GBM cell lines. Importantly, we used the small-molecule non-competitive TrkB antagonist ANA-12 (Cazorla et al., 2011) as a comparison compound in this analysis because it has been reported to reduce A-172 cell viability in a dose- and time-dependent manner (Pinheiro et al., 2017). Hence, this represents a relevant reference measure that also connects with our recent publication demonstrating the interference of cannflavins A and B on TrkB signaling in mouse primary cortical neurons (Holborn et al., 2023).

As presented in **Figure 1**, applying cannflavin A and cannflavin B in the 1-20 µM final concentration range to the culture media of either A-172 (**Fig. 1A-C**) and U-87 (**Fig. 1D-F**) cells produced similar mixed results. Specifically, no significant effect on viability was detected using a resazurin reduction assay for cannflavin A at any concentration or time-point, although a trend for reduced viability can be observed in A-172 cells with the highest concentration (20 µM) at the 24-hour and 48-hour time-points. On the other hand, cannflavin B produced a robust reduction in viability for both A-172 and U-87 cells, with the most significant effect observed at higher concentrations (20 µM) and longer treatment times (48 hours). Interestingly, ANA-12 in the 20-50 µM range also negatively affected survival in both cell lines, with the notable difference that higher compound concentrations were required to achieve an impact comparable to that of cannflavin B. This can be best appreciated at 48 hours in A-172 cells, where 5 µM cannflavin B reduced viability below 50%, while ANA-12 at 10 µM had no statistically significant effect (estimated average cell viability of 90%). The estimated IC50 for each of the compounds and time-points in this experiment are presented in the **Supplementary Information**. Together, these results establish that cannflavin B is more potent at affecting GBM cell survival than cannflavin A and ANA-12.

**Figure 1.**
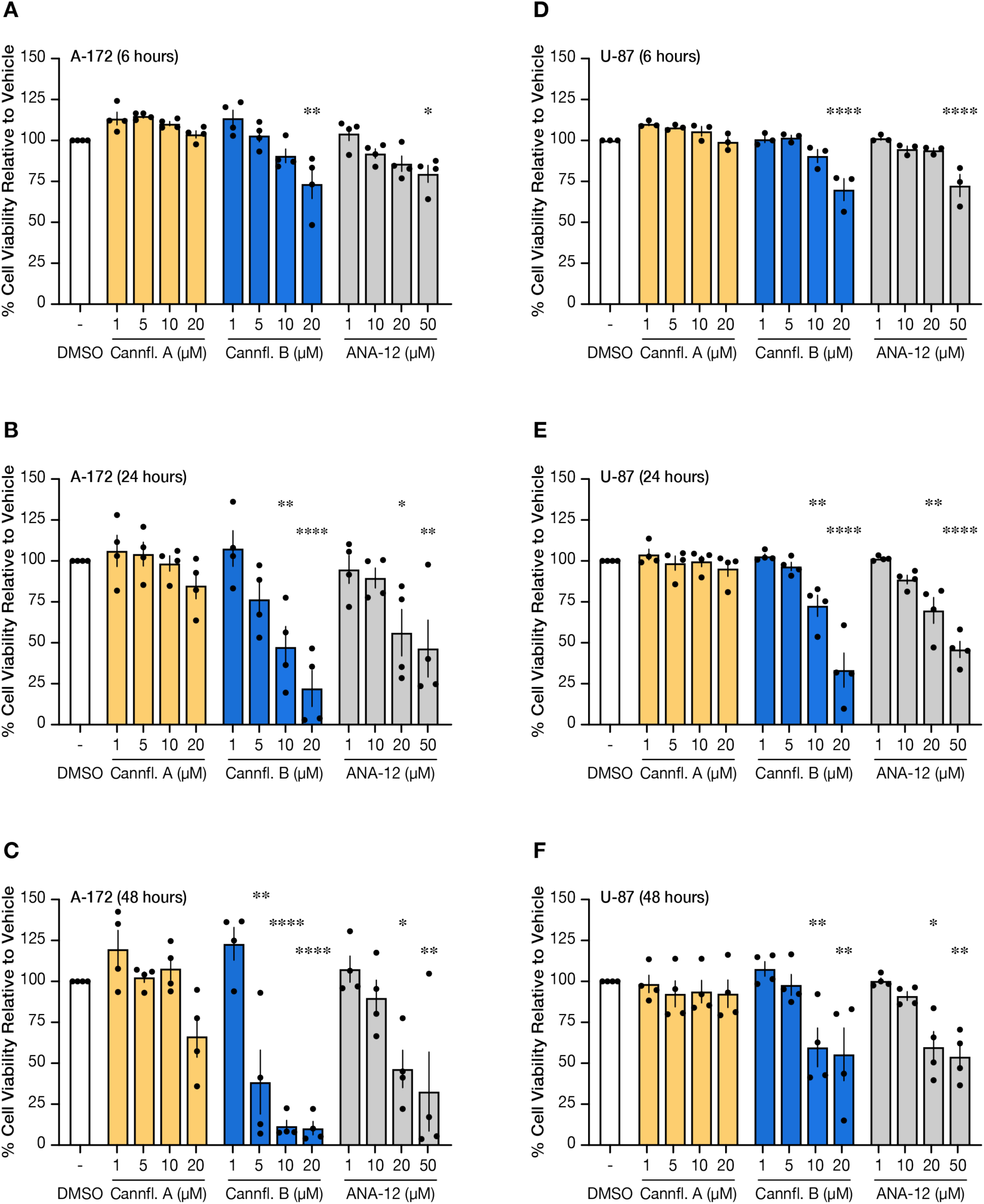
Cannflavin B reduces GBM cell viability. A172 **(A-C)** and U-87 **(D-F)** cells were treated with increased concentrations of cannflavin A (1-20 µM), cannflavin B (1-20 µM), or ANA-12 (1-50 µM) and cell viability was measured at 24-, 48-, and 72-hour time-points using the resazurin reduction assay. DMSO was used as a vehicle control. Application of cannflavin B and ANA-12, but not cannflavin A, reduced the viability of both A-172 and U-87 GBM cells in a dose- and time-dependent manner. A one-way ANOVA was used to analyze the data from each cell line, treatment, and time point independently with DMSO as the reference control. Tukey’s honestly significant difference (HSD) post hoc test: * *p* < 0.05, ** *p* < 0.01, *** *p* < 0.001, **** *p* < 0.0001. Graphs represent mean ± SEM (*n* = 3-4).

To gain a broader view of the potency of cannflavin B towards U-87 cells, we next performed a similar experiment with two structurally related flavones, namely chrysoeriol, the non-prenylated synthesis precursor of cannflavins in the *C. sativa* plant (**Fig. S2A**, Rea et al., 2019), and techtochrysin (**Fig. S2B**), an O-methylated flavone previously discovered to have effects against SW480 and HCT116 colon cancer cells through inhibition of Nuclear factor-κB (NF-κB) activity (Park et al., 2015). Interestingly, we found that the direct application of chrysoeriol to the cell culture medium significantly reduced U-87 cell viability, but only at a final concentration of 20 µM and for treatment times of 24-48 hours (**Fig. S2C**). This result suggests that the prenyl group found at the 3’ position of the flavone B-ring that distinguishes cannflavin B from chrysoeriol contributes to the potency of the latter molecule in this assay. The effect of techtochrysin, however, was the opposite of that of chrysoeriol. Intriguingly, application of techtochrysin in the range of 5-40 µM for 24-48 hours significantly enhanced the viability of U-87 cells. In summary, these experiments reinforce the notion that cannflavin B exerts anti-survival effects on GBM cells at lower concentrations and more rapidly than closely related flavones.

### Cannflavins A and B do not induce necrosis in GBM cells

To assess the extent to which cannflavin B’s observed impact on GBM cell viability could be attributed to necrosis, we performed cell membrane integrity tests with the lactate dehydrogenase (LDH) release assay. A-172 and U-87 cells treated with 10-20 µM cannflavin A or cannflavin B for 24 hours presented minimal LDH release (**Fig. S3A, B**). Specifically, application of cannflavin A resulted in 3.57% LDH release at 10 µM and 2.54% at 20 µM in A-172 cells, and 0.96% at 10 µM and 1.03% at 20 µM in U-87 cells. For cannflavin B, LDH release was 4.43% at 10 µM and 5.38% at 20 µM in A-172 cells, and 0.81% at 10 µM and 0.88% at 20 µM in U-87 cells. Those values are comparable to those obtained with ANA-12 at 10 µM (2.01% for A-172 cells and 1.61% for U-87). In summary, these results suggest that the adverse effect of cannflavin B on cell viability, as assessed using the resazurin reduction method, is not due to necrotic cell death at the specified concentrations and time points.

### Cannflavins A and B do not cause apoptosis in the low micromolar range

Our following goals were to assess the effect of cannflavins A and B in the low micromolar (1 to 10 µM) range on GBM’s cell cycle, migration, and capacity to invade. Before proceeding with those tests, though, we wanted to confirm that neither cannflavin A nor cannflavin B causes widespread programmed cell death resulting in DNA fragmentation after at least 24 hours of exposure. Using the highly sensitive TUNEL assay, we observed that A-172 GBM cells treated with cannflavin A or cannflavin B at a final concentration of 10 µM for 24 hours did not produce a noticeable fluorescein signal indicative of apoptosis (**Fig. S3C**). This negative result assures that any effect associated with cannflavins at a concentration of 10 µM or lower, which will be identified next, is distinct from programmed cell death.

### Effect of cannflavins on A-172 and U-87 cell cycle

Various flavonoids have been demonstrated to affect the cell cycle of different cancer cell types (e.g., Lin et al., 2015; Srivastava et al., 2016; Zhang et al., 2017). This led us to ask whether cannflavins A and B, at concentrations that do not produce a rapid and significant anti-viability effect on A-172 and U-87 cells, could alter their cell cycle. Flow cytometry analysis showed that neither cannflavin A nor cannflavin B affected cell cycle phase after 24 hours of treatment in the concentration range of 1-5 µM (**Fig. 2**). Interestingly, the A-172 and U-87 cell lines exhibited distinctive distributions of the cell cycle that remained unchanged with treatment (G1 proportion for the DMSO control was 61.7% and 41.6%, respectively). In both GBM cell lines, compared to the DMSO control, there was no statistically significant change in the cell cycle phase upon cannflavin A or cannflavin B treatment. These results demonstrate that cannflavins in the low micromolar range (1 to 10 µM) do not alter GBM cell cycle.

**Figure 2.**
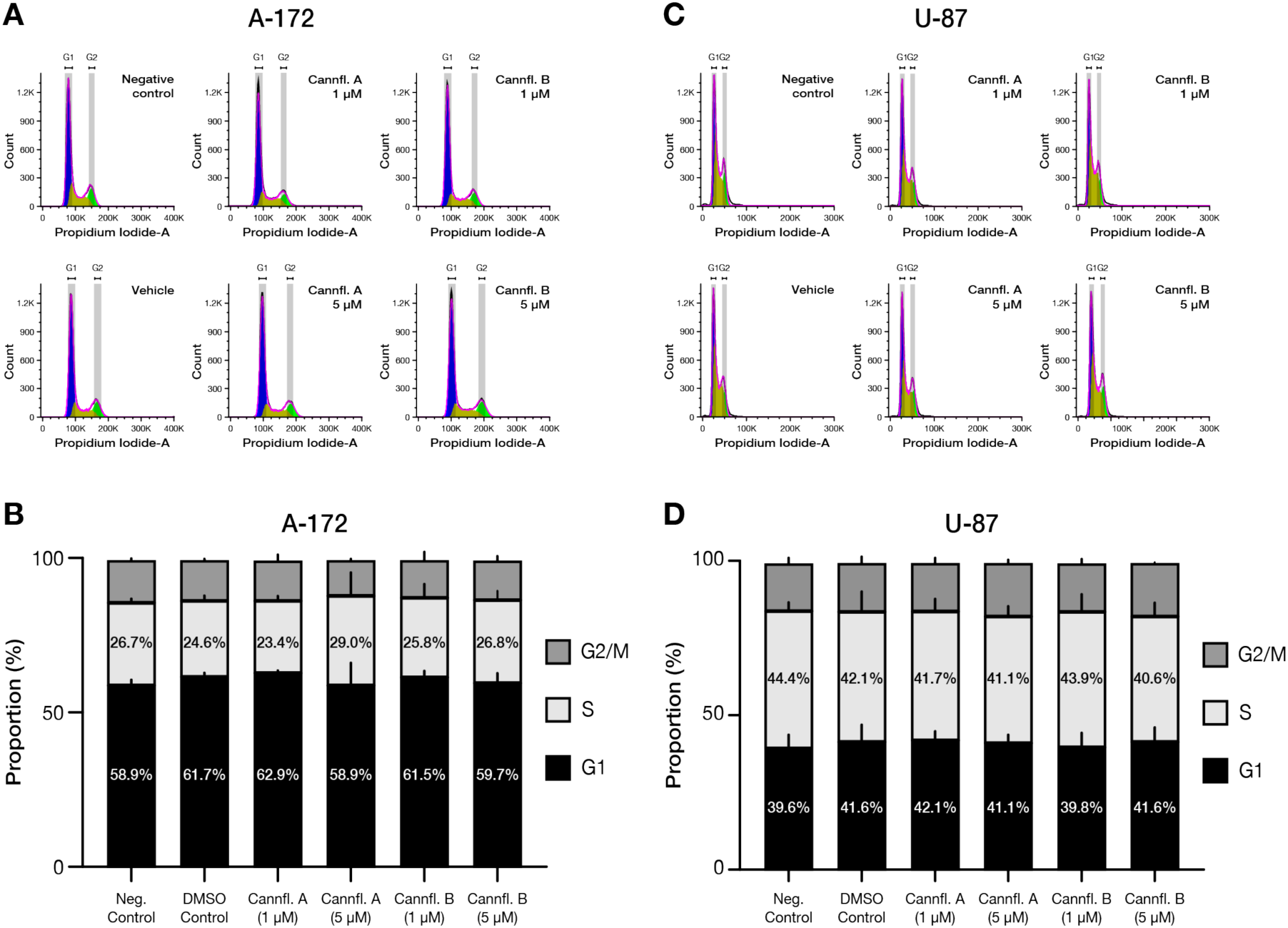
Cannflavins A and B did not affect cell cycle distribution in GBM cell lines. A-172 **(A-B)** and U-87 **(C-D)** DNA content was determined by gating on the G1, S, and G2/M phases using flow cytometry and analyzed using the Watson Pragmatic algorithm to distribute cell cycle phases. A172 **(B)** and U-87 **(D)** cell cycle data are expressed as mean ± SEM (*n* = 3).

### Cannflavins limit GBM cell migration and invasion

Invasion and metastasis are primarily responsible for mortality associated with solid tumors (Li et al., 2025). This is the case for GBM, where rapid invasion and damaging infiltration of the cancer cells into surrounding healthy brain tissue account for most of the poor outcomes (Barbaro et al., 2022). Therefore, systematically assessing the action of cannflavins on GBM cell migration and invasion is highly relevant.

To start, we completed an analysis of cell migration using a time-lapse scratch assay in which cannflavin A (10 µM final concentration), cannflavin B (5 µM final concentration), ANA-12 (10 µM final concentration), or DMSO alone was applied directly to the culture media of A-172 cells. Cell migration into the scratch wound area was monitored over 48 hours (**Fig. 3A** and **Supplementary Video**). Our data revealed striking differences between wells treated with cannflavins and exposed to DMSO. First, analysis of the binary area fraction over all time-points suggested a delayed wound closure in inhibitor-treated conditions and more linear cell movement over time for the DMSO and cannflavin A conditions compared to the cannflavin B and ANA-12 (**Fig. 3B**). Examining next just the 24-hour and 48-hour time-points, we found that the normalized wound area fraction was significantly lower for the three tested molecules compared to DMSO (**Fig. 3C**). Further, considering the data as a percentage of wound closure, significantly less area was covered by cells after 24 hours and 48 hours exposure to cannflavin B, but only at 48 hours for cannflavin A (**Fig. 3D**). ANA-12 results in that analysis were not different compared to DMSO. Finally, considering the number of individual cells detectable within the wound area, significantly fewer were present at the end of the experiment for cannflavin B (**Fig. 3E**).

**Figure 3.**
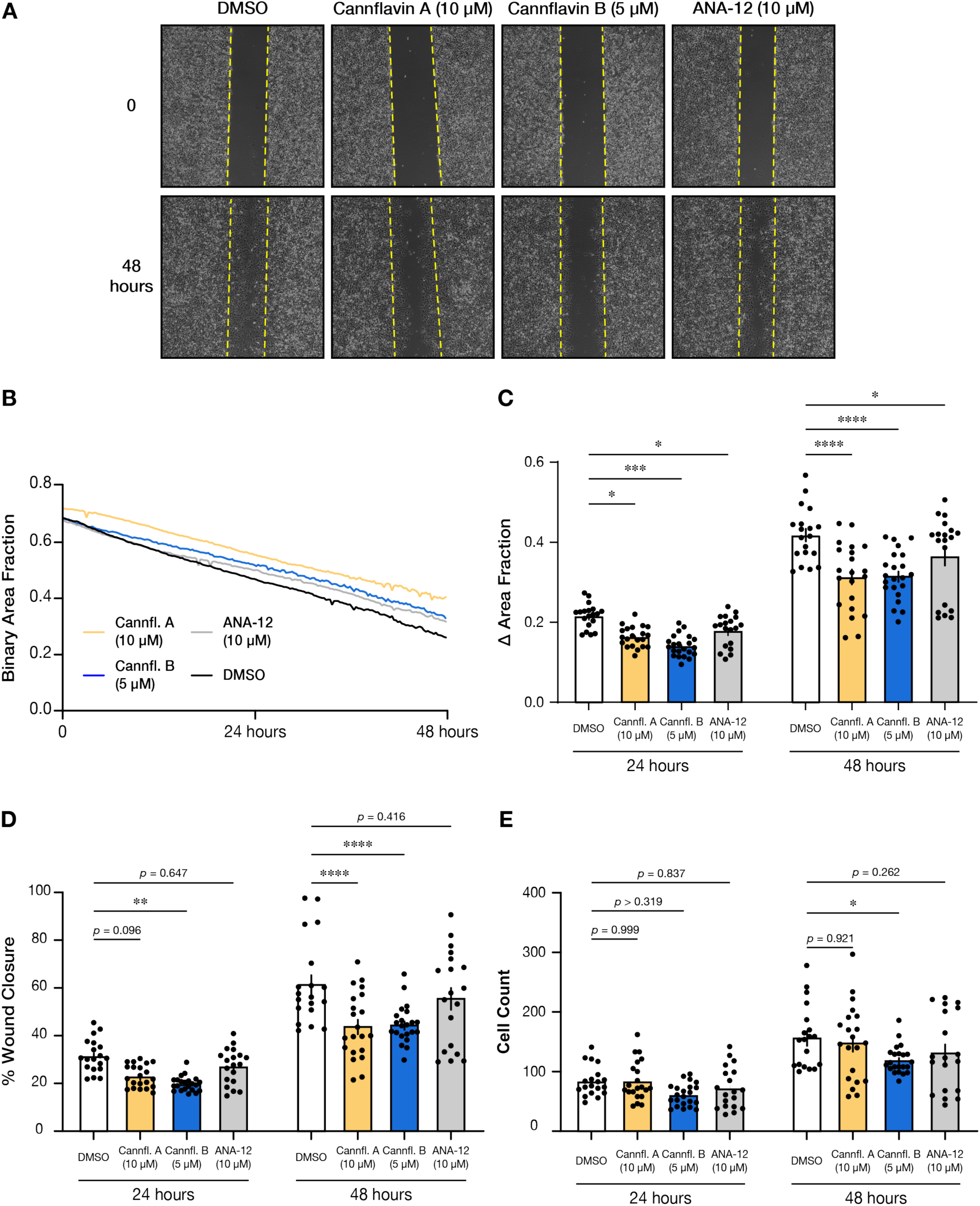
Cannflavins A and B suppress A-172 cell migration. **(A)** Representative phase-contrast images of scratch assay showing A-172 GBM cancer cells within a gap created with a pipette tip at 0 hour and 48 hours. Dashed yellow lines indicate the initial wound edges. **(B)** Quantification of the binary wound area fraction over time, showing delayed wound closure in inhibitor-treated conditions. **(C)** Normalized wound area fraction at 24 hours and 48 hours. A two-way ANOVA reveals a significant effect of time (*F*_1,218_ = 133.3, *p* < 0.0001) as well as treatment (*F*_3,218_ = 18.37, *p* < 0.0001). There is no significant interaction between time and treatment (*F*_3,218_ = 2.094, *p* = 0.1019). Post hoc comparisons between DMSO control and each treatment means were conducted using Dunnett’s multiple comparisons. All treatments at all time points produced a significant reduction in cell migration. **(D)** Percentage of wound closure at 24 hours and 48 hours. A two-way ANOVA reveals a significant effect of time (*F*_1,218_ = 104.2, *p* < 0.0001) as well as treatment (*F*_3,218_ = 16.45, *p* < 0.0001). There is no significant interaction between time and treatment (*F*_3,218_ = 1.885, *p* = 0.133). Post hoc comparisons between DMSO control and each treatment means were conducted using Dunnett’s multiple comparisons. Cannflavin B significantly limited closure at both time points, while cannflavin A was significant only at 48 hours. ANA-12 did not cause a significant effect. **(E)** Total cell counts within the imaging field at 24 hours and 48 hours. A two-way ANOVA reveals a significant effect of time (*F*_1,154_ = 92.25, *p* < 0.0001) as well as treatment (*F*_3,154_ = 4.46, *p* = 0.0049). There is no significant interaction between time and treatment (*F*_3,154_ = 0.2556, *p* = 0.8573). Post hoc comparisons between DMSO control and each treatment means were conducted using Dunnett’s multiple comparisons. Significantly less cells were found in the wound area with cannflavin B at 48 hours. No other comparisons were significantly different. All data represented as mean ± SEM (3 biological replicates with 6-8 images per well; * *p* < 0.05, ** *p* < 0.01, *** *p* < 0.001, **** *p* < 0.0001).

Next, we performed a transwell assay to gauge the potential of cannflavin B to limit A-172’s capacity to invade when the cells are exposed to the molecule. A-172 cells were pre-treated overnight (∼16 hours) with 10 µM cannflavin B or DMSO and then plated on Matrigel-coated inserts with the addition of fresh cannflavin B (10 µM). As presented in **Figure 4**, the results from the transwell invasion assay revealed that exposure to cannflavin B significantly reduced the number of cells that invaded through the Matrigel matrix, resulting in a statistically significant reduction of more than 50% compared to the vehicle control. This result supports that cannflavin B at a concentration of 10 µM effectively inhibits GBM cell migration and invasion, providing initial evidence of its potential against tumor progression.

**Figure 4.**
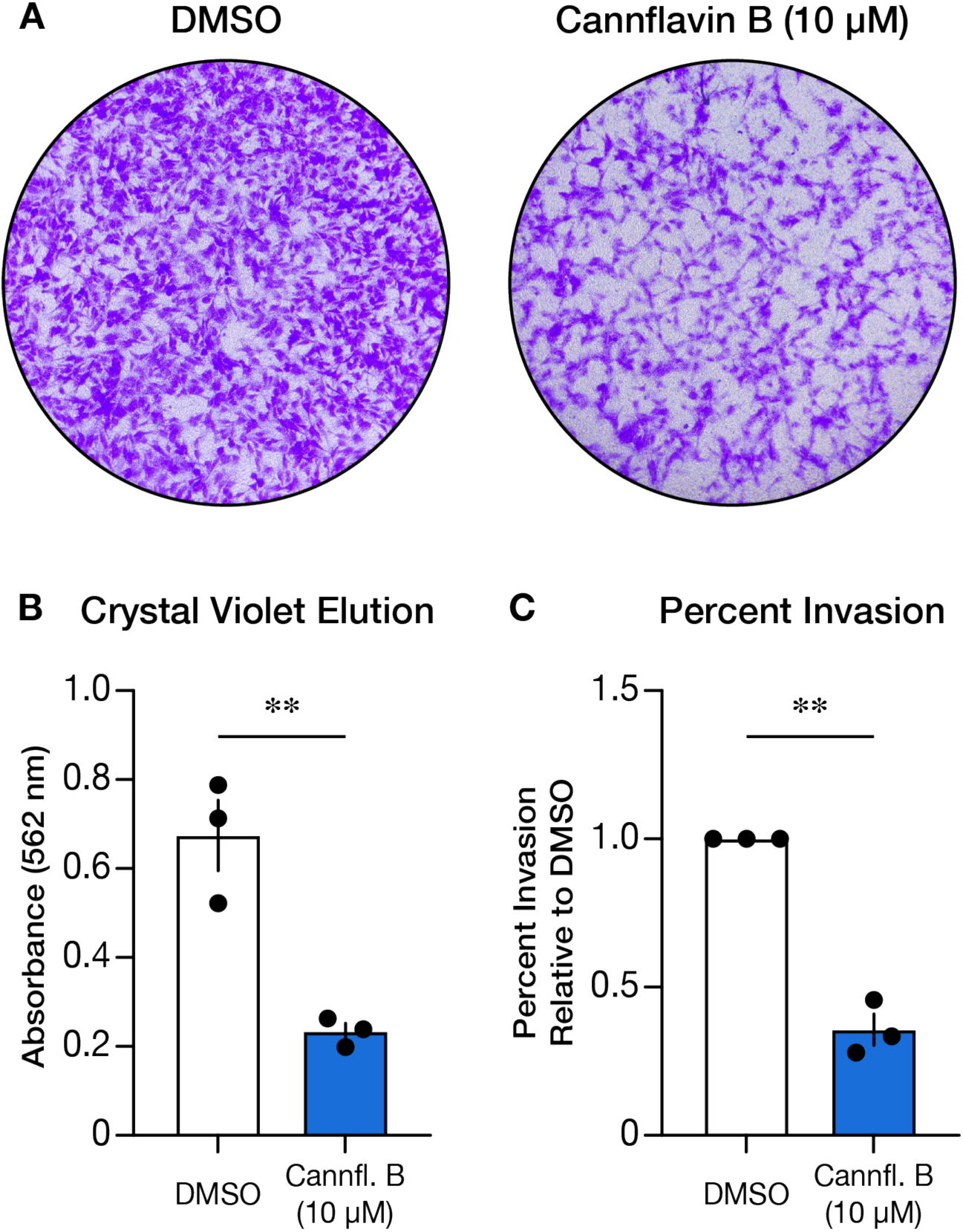
Cannflavin B inhibits A-172 glioblastoma cell invasion in a transwell assay. **(A)** Representative images of stained migrated A-172 cells following treatment with DMSO or cannflavin B (10 µM). **(B)** Quantification of migration using crystal violet elution measured by absorbance at 562 nm. **(C)** Percent invasion between DMSO control and cannflavin B, showing a significant reduction in cannflavin B-treated cells. Data represent mean ± SEM, with statistical significance determined by an unpaired two-tailed *t*-test (** *p* < 0.01).

Lastly, since U-87 cells have a strong tendency to aggregate and form tumor-like spheroid formations (tumorspheres) in suspension culture, we leverage this attribute to provide a different evaluation of the action of cannflavins A and B on the invasion ability of these GBM cell subtypes. Specifically, we cultured U-87 as 3-dimensional (3D) tumorspheres and then embedded those within a Matrigel matrix to measure how they would progress in the outward direction (**Fig. 5A**). Interestingly, adding cannflavin A to the culture media at a final concentration of 10 µM resulted in a significantly smaller radius and area of tumorsphere outgrowth at the three examined times (**Fig. 5B, C**). For cannflavin B, tumorspheres in wells treated at a final concentration of 5 µM presented a significantly reduced area at 72 hours after the start of the experiment (**Fig. 5B**) and a smaller radius in the 48-72 hours times (**Fig. 5C**). ANA-12, for its part, was minimally active in this assay producing only a slight significant effect on radius at 72 hours (**Fig. 5C**). Taken together, these three experiments provide strong evidence that cannflavins, particularly cannflavin B, can produce rapid and sustained anti-migration and anti-invasion action against GBM cells.

**Figure 5.**
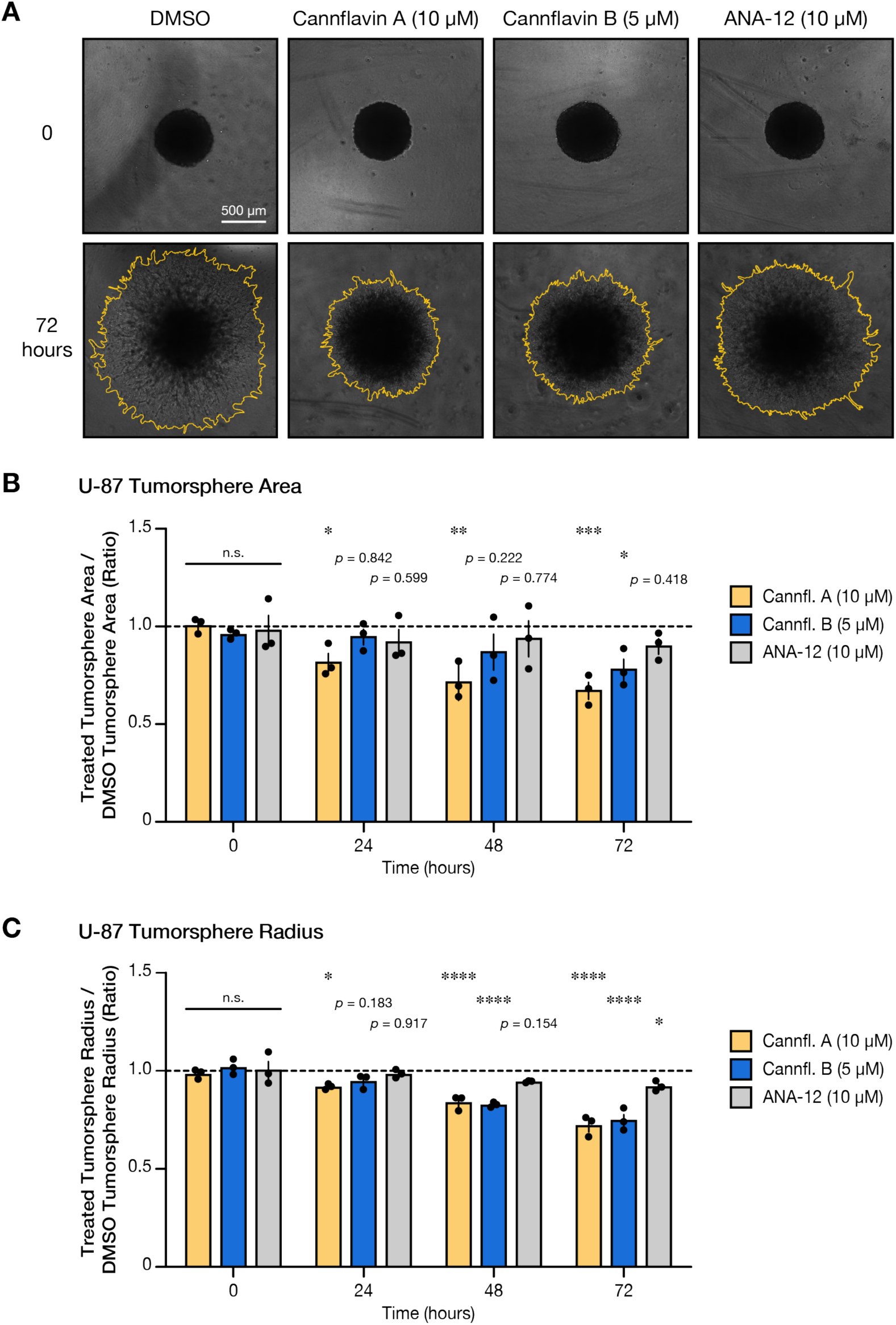
Cannflavins A and B inhibit the outgrowth of U-87 tumorspheres. **(A)** Representative phase-contrast images of U-87 tumorspheres at 0 hour and 72 hours post-treatment with DMSO (vehicle), ANA-12 (10 µM), cannflavin A (10 µM), or cannflavin B (5 µM) in a tumorsphere invasion assay. Scale bar: 500 µm. The white outline indicates the borders of tumorsphere outgrowth. **(B)** Quantification of relative tumorsphere area expansion over time, normalized to the vehicle control. A two-way ANOVA reveals a significant effect of time (*F*_3,32_ = 5.949, *p* < 0.0024) as well as treatment (*F*_3,32_ = 10.13, *p* < 0.0001). There is no significant interaction between time and treatment (*F*_9,32_ = 1.765, *p* = 0.1145). Post hoc comparisons between DMSO control and each treatment means were conducted using Dunnett’s multiple comparisons. Ratio was significantly lower at all three time points for cannflavin A, while cannflavin B was significant only at 72 hours. ANA-12 did not cause a significant effect. **(C)** Relative tumorsphere radius expansion over time, showing significant inhibition of invasion upon cannflavin treatment. A two-way ANOVA reveals a significant effect of time (*F*_3,32_ = 46.18, *p* < 0.0001) as well as treatment (*F*_3,32_ = 40.07, *p* < 0.0001). Notably, there is a significant interaction between time and treatment (*F*_9,32_ = 8.749, *p* < 0.0001), reflecting an earlier effect with cannflavin A, followed by cannflavin B, and finally ANA-12. Post hoc comparisons between DMSO control and each treatment means were conducted using Dunnett’s multiple comparisons. Data represent mean ± SEM (*n* = 3; * *p* < 0.05; ** *p* < 0.01; *** *p* < 0.001; **** *p* < 0.0001; n.s., not significant).

### Cannflavin B limits the growth and infiltration of U-87 GBM tumorspheres attached to cerebral organoids

Conventional pre-clinical studies for GBM consist of monolayer tumor cell cultures and animal approaches, including xenografts and genetically engineered mouse models (Gómez-Oliva et al., 2021; Haddad et al., 2021). However, these strategies encounter significant obstacles in recapitulating the complexity of GBM, especially the variety of tumor cellular states and the cancerous mass microenvironment. The use of brain organoids derived from human pluripotent stem cells helps overcome some of those challenges and provides exciting new ways to study GBM tumor formation and invasion in real time (Linkous et al., 2019; Wang et al., 2023)—an approach that our group has used with success in recent years (Tilak et al., 2021b; Lau et al., 2025). To test if cannflavin B could produce therapeutically relevant effects in a more complex *in vitro* system, we conducted a co-culture experiment where iPSC-derived cerebral organoids matured for 60 days were mixed with U-87 tumorspheres expressing GFP (**Fig. 6A**). Our pool of co-culture specimens was then divided in two groups: one where the U-87 tumorspheres attached and grew unchallenged for 11 days followed by the application of cannflavin B at a final concentration of 5 µM in the culture media for 3 days (Paradigm 1), and another (Paradigm 2) where the cancer cells attached and grew for 4 days and then 5 µM cannflavin B was added for 10 days (**Fig. 6B**). Co-cultures treated with DMSO only were included for both tests as control, and media change with fresh cannflavin B (5 µM) or DMSO was done every other day for the duration of the experiment. As seen in fluorescence microscopy captures of immunostained sections (**Figure 6C-F**), U-87 tumorspheres attached to cerebral organoids in both paradigms. In the two DMSO control conditions (**Fig. 6C, E**), a distinct intermingling between GBM cells expressing GFP and neurons, as revealed by TUJ1 immunostaining, can be observed at the edge of the cerebral organoid where the tumorsphere is attached. Interestingly, less GBM cell penetration into the organoids was observed in the cannflavin B-treated wells in both tests (**Fig. 6D, F**). To quantify this phenotype, we determined the proportion of pixels corresponding to the GFP signal (U-87 GBM cells) in their respective cerebral organoid section (**Fig. 6G, H**). Although this analysis did not reveal statistical significance due to low GBM cell infiltration in some DMSO control replicates, cannflavin B-treated samples consistently showed very little GFP signal. Considering this fact, we consider this result highly consistent with other findings demonstrating the efficacy of cannflavin B against GBM cells. Finally, although the tumorsphere total area for corresponding sections was not statistically different between the two conditions in the 3-day (Paradigm 1) experiment (**Fig. 6I**), we found that the U-87 tumorspheres were significantly smaller when cannflavin B was added for 10 days (Paradigm 2, **Fig. 6J**). Together, these observations collected with a more physiologically relevant model provide compelling additional data to support an effect of cannflavin B against the ability of aggressive GBM cells to invade human brain-like tissue.

**Figure 6.**
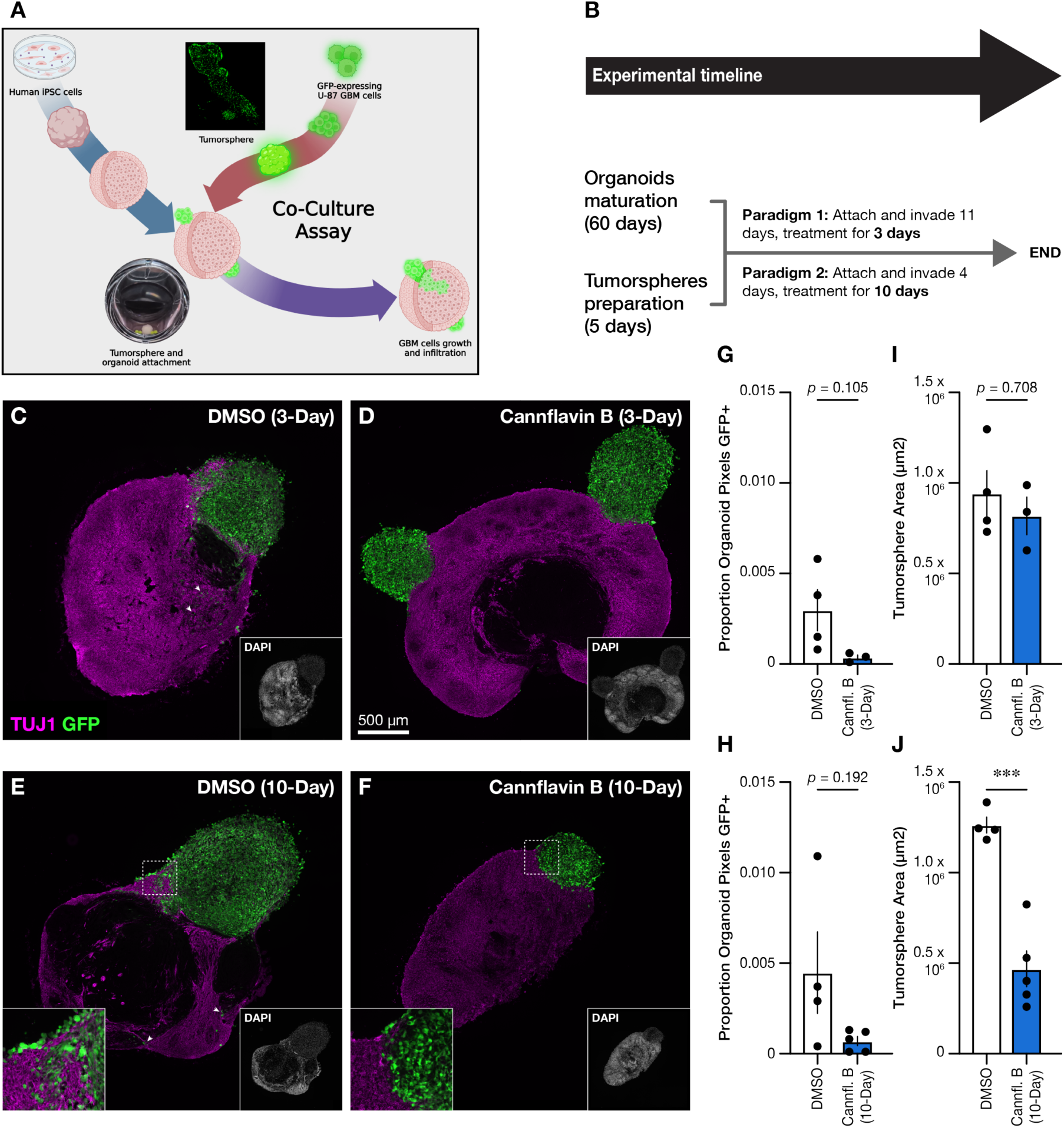
Cannflavin B reduces the size of attached tumorspheres to a cerebral organoid. **(A)** Schematic representation of cerebral organoid and U-87 tumorspheres co-culture model. Cerebral organoids matured for 60 days before introducing tumorspheres prepared with U-87 cells expressing GFP. **(B)** Two experimental timelines were tested: 11 days of co-culture followed by 3 days of applying cannflavin B to the culture media (final concentration of 5 µM), and 4 days of co-culture followed by 10 days of applying cannflavin B (final concentration of 5 µM). **(C-F)** Representative capture of organoid with attached tumorsphere(s) for all conditions. **(G-H)** Proportion of GFP-positive pixels corresponding to U-87 cells within the cerebral organoid section area for the 3-day **(G)** and 10-day **(H)** conditions. **(I-J)** Estimation of tumorsphere size in the 3-day **(I)** and 10-day **(J)** conditions. The average tumorsphere area (GFP-positive cells, green) was significantly smaller after 10 days of cannflavin B treatment than in the corresponding DMSO control. Data represent mean ± SEM, with statistical significance determined by an unpaired two-tailed *t*-test (*** *p* < 0.001).

### Cannflavins A and B impact on TrkB signaling in GBM cells

As mentioned previously, a study by Pinheiro and colleagues (2017) reported that the TrkB inhibitor ANA-12 reduces the viability of A-172 cells, suggesting that the activity of this RTK may be essential for the growth and survival of this GBM cell line. Since our previous work using mouse primary cortical neurons also found that cannflavins A and B can interfere with TrkB signaling (Holborn et al., 2023), we considered it necessary to gain more information about the effect of cannflavins and ANA-12 on TrkB signaling in A-172 (**Fig. 7**) and U-87 (**Fig. 8**) cells to interpret our functional data. As expected, western blot analysis revealed that application of cannflavin A at a final concentration of 20 µM (**Fig. 7A, B**) and cannflavin B at 10-20 µM (**Fig. 7F, G**) to the culture media of A-172 cells for 6 hours significantly reduced the phosphorylation of TrkB at tyrosine 705 (Y705), which is in the catalytic domain and stimulates kinase activity of the receptor when phosphorylated. Consistent with this result, ANA-12 at a final concentration of 10 µM also significantly decreased phospho-TrkB Y705 signal (**Fig. 7B, G**). Interestingly, however, our western blot data also revealed two unexpected and critical results. Firstly, key downstream TrkB signaling effectors (i.e., p44/p42 MAPK and AKT) were unaffected by cannflavins and ANA-12 in A-172 cells (**Fig. 7C-E, H-J**). Secondly, cannflavins and ANA-12 did not impair TrkB phosphorylation and downstream signaling effectors in U-87 cells, even at the highest tested concentration (**Fig. 8**). Together, these western blot data force the critical conclusion that any positive results about GBM survival, migration, and invasion presented above are most likely distinct from TrkB signaling impairment, whether they were the result of cannflavins or ANA-12 treatment.

**Figure 7.**
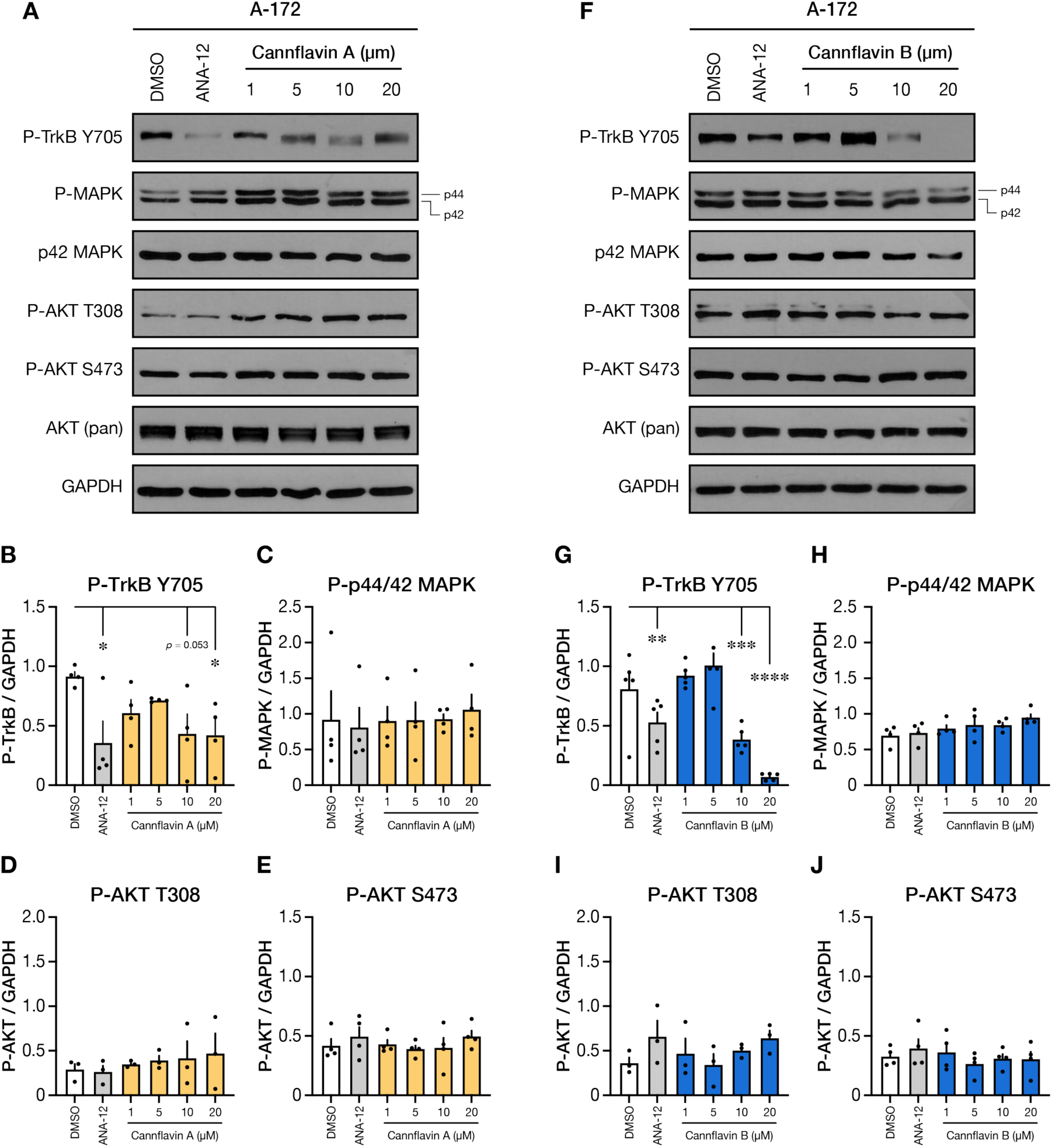
Cannflavin A and cannflavin B affect TrkB Y705 activation site but not downstream signaling in A-172 cells. Western blot analysis from A-172 cells treated for 6 hours with cannflavin A **(A-E)** and cannflavin B **(F-J)** at various concentrations. Corresponding densitometry quantifications reveal a significant reduction in P-TrkB Y705 signal, but not in P-MAPK and P-AKT. GAPDH (glyceraldehyde-3-phosphate dehydrogenase) was used as a loading control. Graphs show the mean ± SEM (*n* = 4) of the target/GAPDH ratio for cells treated as in **(A)** and **(F)**. Post hoc comparisons between DMSO control and each treatment means were conducted using Dunnett’s multiple comparisons (* *p* < 0.05; ** *p* < 0.01; *** *p* < 0.001; **** *p* < 0.0001).

**Figure 8.**
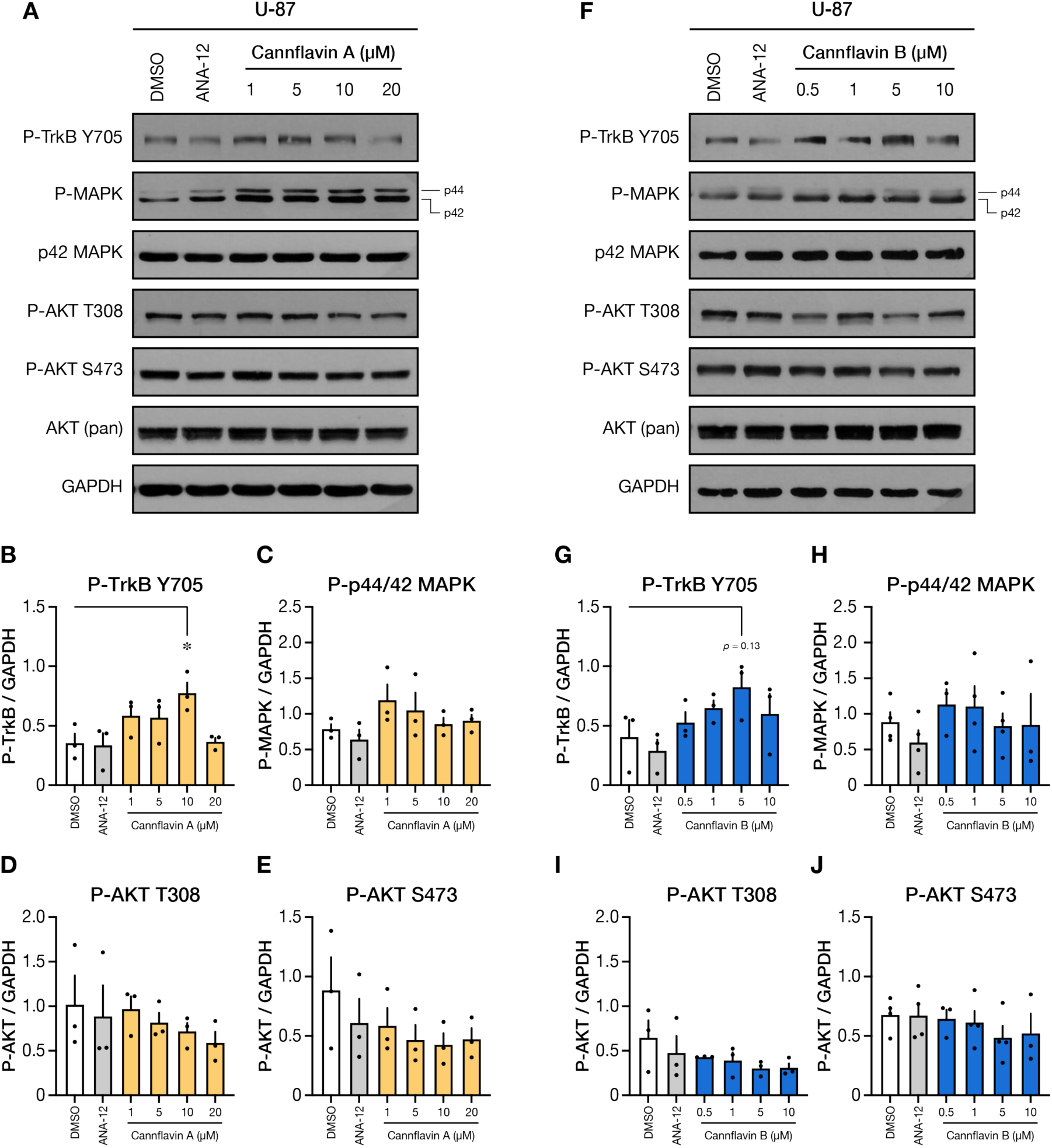
Cannflavin A and cannflavin B slightly increase TrkB Y705 activation site but not change downstream signaling in U-87 cells. Western blot analysis from U-87 cells treated for 6 hours with cannflavin A **(A-E)** and cannflavin B **(F-J)** at various concentrations. Corresponding densitometry quantifications reveal a significant reduction in P-TrkB Y705 signal, but not in P-MAPK and P-AKT. GAPDH (glyceraldehyde-3-phosphate dehydrogenase) was used as a loading control. Graphs show the mean ± SEM (*n* = 3) of the target/GAPDH ratio for cells treated as in **(A)** and **(F)**. Post hoc comparisons between DMSO control and each treatment means were conducted using Dunnett’s multiple comparisons (* *p* < 0.05).

## Discussion

In this study, we employed various *in vitro* assays to characterize the phenotypic outcomes of cannflavins A and B on the viability, cell cycle, migration, and invasion capacities of A-172 and U-87 GBM cells. Of particular interest, our results show a consistent dose-dependent decrease in cell viability in both lines when cannflavin B is added to the culture media. No effect on cell cycle by either cannflavin molecule was detected, but we found that both molecules can reliably limit tumor cell migration below the 10 µM concentration and that cannflavin B has robust anti-invasive properties based on results from transwell and tumorsphere assays. Together, these observations underscore the multifaceted therapeutic potential of cannflavin B against this aggressive type of brain cancer.

Focusing first on our overall cell survival results, it is interesting to speculate about why cannflavin B shows a higher effect against A-172 and U-87 cells than the other tested flavonoids (i.e., cannflavin A, chrysoeriol, and techtochrysin). One key factor contributing to this effect is certainly prenylation, which is considered to increase the lipophilicity of flavonoids, thereby enhancing their affinity for biological membranes and improving their interactions with target proteins (Xu et al., 2012; Yang et al., 2015). The reduced potency of chrysoeriol, which is structurally similar to cannflavins A and B but lacks prenylation attached to the B-ring (**Fig. S2A, C**), is consistent with this interpretation. That said, the length and position of the prenyl side-chain are undoubtedly also critical, as we measured greater bioactivity with cannflavin B, which has a single prenyl group, compared to cannflavin A, which has two (**Fig. S1**). Finally, another structural-activity feature that may explain other differences among flavonoids in our study is the B-ring hydroxylation patterns. In fact, extensive evidence suggests that hydroxylation at specific positions can markedly enhance bioactivity (Liu et al., 2022; Li et al., 2023). This detail most likely contributed to the result we obtained in the techtochrysin test (Fig. S2B, D).

In addition, we conducted a western blot analysis to gain insights about the possible molecular signaling changes induced by cannflavins that could produce the reported cellular effects. However, the tested cannflavins and ANA-12 consistently reduced TrkB phosphorylation at Y705 in A-172 cells, but this effect was not observed in U-87 cells. The divergent TrkB phosphorylation responses observed in A-172 and U-87 cells likely reflect differences in baseline endogenous TrkB expression (**Fig. S4**) and downstream pathway coupling. U-87 cells are known to express lower levels of full-length TrkB than A-172 cells, which may explain their limited responsiveness. Most importantly, cannflavins and ANA-12 had minimal effects on MAPK and AKT, which are well-known contributing factors to cancer hallmarks, including growth, apoptosis, and invasion. These unexpected results depart from our previous work on cannflavins A and B in mouse primary cortical neurons (Holborn et al., 2023) and suggest that different molecular events, influenced by cannflavins, are responsible for the migration and invasion disruptions observed in GBM cells in this study. Therefore, future work is needed to elucidate the target engagement of cannflavins in GBM cells. One unbiased strategy for this task could be to utilize the Proteome Integral Solubility Alteration (PISA) assay, a high-throughput and mass-spectrometry-based approach that leverages binding-dependent changes in protein solubility to identify drug-target interactions (Gaetani et al., 2019; Gaetani and Zubarev, 2023).

On a separate note, an intriguing finding in our study concerns the increased viability of U-87 cells exposed to techtochrysin (**Fig. S2B, D**)—a flavone molecule that had been reported to suppress the growth of SW480 and HCT116 human colon cancer cells by inhibiting NF-κB resulting in the expression of death receptors like DR3, DR4, and Fas (Park et al., 2015). Although we cannot explain at this point why U-87 cells exposed to this specific flavone produced an opposite effect to what has been reported with human colon cancer cells, this bring to our attention the possibility that some of the anti-cancer phenotypes associated with cannflavins A and B that we found may be linked to disruption of the transcription factor NF-κB. Indeed, extensive research has implicated NF-κB in various aspects of cancer formation, growth, treatment response, immune escape, and other related processes (Taniguchi and Karin, 2018; Zhang et al., 2021; Khan et al., 2024). In GBM and other gliomas, it has been reported for more than two decades that the aberrant constitutive activity of this transcription factor contributes to disease initiation and progression, including the invasion phenotype (Nagaai et al., 2002; Raychaudhuri et al., 2004; Wang et al., 2004). Considering this, one tantalizing scenario is the disruption by cannflavins of the constitutive signaling pathway that phosphorylates and promotes degradation of IκBɑ by the ubiquitin-proteasome system, ultimately releasing NF-κB dimers that translocate to the nucleus and modulate target gene expression that contributes to disease progression (Cahill et al., 2016). While the present study did not directly evaluate NF-κB activity, these findings raise the possibility that cannflavin B may influence NF-κB–associated invasion programs. Future work incorporating NF-κB reporter assays and transcriptomic profiling will be essential to define the signaling changes underlying the anti-invasive effects observed here. If this is the case, it would be revealing to test the effect of cannflavin B administration in xenograft or induced mouse models of GBM as used in previous work targeting aberrant NF-κB activity (Friedmann-Morvinski et al., 2016). Ultimately, this would provide a direct test of this hypothesis and additional pre-clinical evidence to promote cannflavin B as a potentially valuable therapeutic agent against GBM.

Lastly, we present evidence that GFP-expressing U-87 cells migrating from a tumorsphere can be challenged by cannflavin B at low µM levels as they infiltrate human iPSC-derived cerebral organoids. Although we could not obtain precise quantitative measures of GBM cell infiltration into the cerebral organoid structure using thin frozen sections, we found that the tumorsphere size was significantly smaller after 10 days of exposure to cannflavin B than after exposure to DMSO (vehicle control). This result calls for a deeper investigation into the therapeutic value of cannflavin B against GBM or the brain metastasis of other cancer types using live animal approaches, as mentioned above. Corroborating the effectiveness of cannflavin B at limiting cancer progression with future animal models would advance our research towards the design and proposal of a clinical trial with human patients.

In conclusion, the distinctive ability of cannflavins, particularly cannflavin B, to inhibit GBM cell migration and invasion, combined with their restrained cytotoxicity, positions them as promising candidates for developing novel therapeutic strategies against GBM (**Fig. S5**). Future studies should focus on validating these effects in animal models to understand the pharmacokinetics, bioavailability, and therapeutic index of cannflavins. Furthermore, exploring the synergistic impact of cannflavins with existing chemotherapeutic agents or targeted therapies could provide valuable insights into their potential as adjuvant treatments, particularly in overcoming drug resistance—a significant challenge in the management of GBM. Combining cannflavins with agents like TMZ (Stupp et al., 2005; Stupp et al., 2009; Fisher et al., 2013) or the anti-angiogenic agent bevacizumab (Fu et al., 2023) could enhance their therapeutic efficacy by simultaneously targeting multiple aspects of tumor growth and invasion.

## Abbreviations

AKT: Protein kinase B
BDNF: Brain-Derived Neurotrophic Factor
BSA: Bovine Serum Albumin
DMEM: Dulbecco’s Modified Eagle’s Medium
DMSO: Dimethylsulfoxide
FBS: Fetal Bovine Serum
GBM: Glioblastoma multiform
GFP: Green Fluorescent Protein
iPSC: induced Pluripotent Stem Cell
MAPK: Mitogen-Activated Protein Kinase
mTOR: mammalian target of rapamycin
NF-κB: Nuclear factor-κB
PBS: Phosphate Buffered Saline
RTK: Receptor Tyrosine Kinase
TMZ: Temozolomide
TrkB: Tropomyosin receptor kinase B

## Data availability

All data collected for the study are available upon inquiry to the corresponding author.

## Consent for publication

Consent to publish the article has been obtained from all authors.

## CRediT Authorship Contribution Statement

**Jennifer Holborn**: Conceptualization, Data curation, Formal analysis, Funding acquisition, Investigation, Writing – original draft, **Hannah Robeson:** Data curation, Methodology, Writing – review & editing. **Ellis Chartley:** Formal analysis, Methodology, Writing – review & editing. **Tiana Gluscevic**: Data curation, Methodology. **Adina Borenstein:** Methodology. **Colby Perrin:** Methodology. **Begüm Alural:** Methodology. **Himain Perera:** Visualization, Writing – review & editing. **Tariq A. Akhtar:** Resources. **Nina Jones:** Resources, Writing – review & editing. **Jasmin Lalonde:** Conceptualization, Data curation, Funding acquisition, Supervision, Writing – original draft.

## Acknowledgments

We wish to thank our lab colleagues for their help with this project.

https://www.editorialmanager.com/PHYPLU/.

## Supplementary Information

**Supplementary Figure 1.**
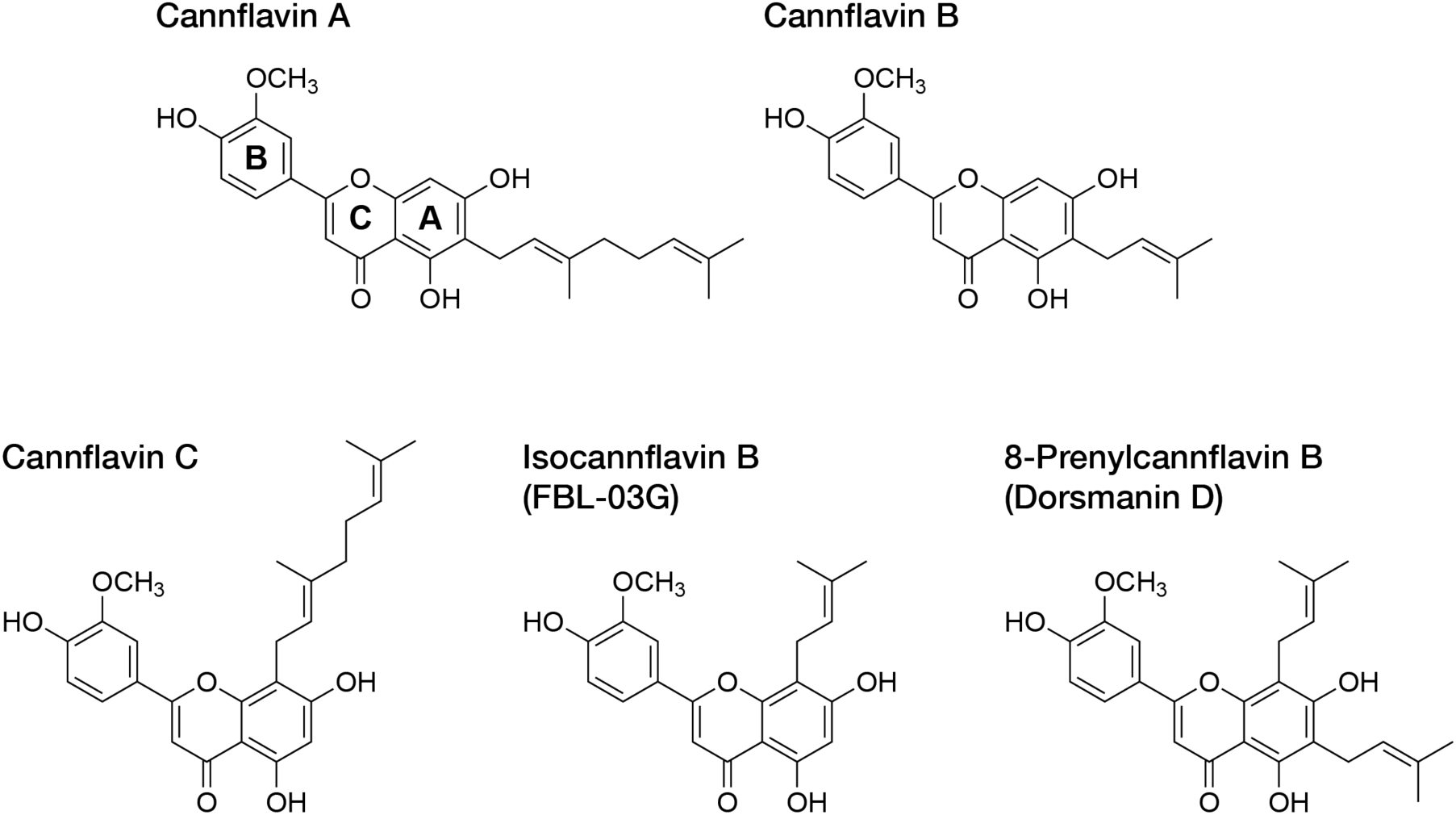
Members of the cannflavin family. Position of each ring is labeled for cannflavin A.

**Supplementary Figure 2.**
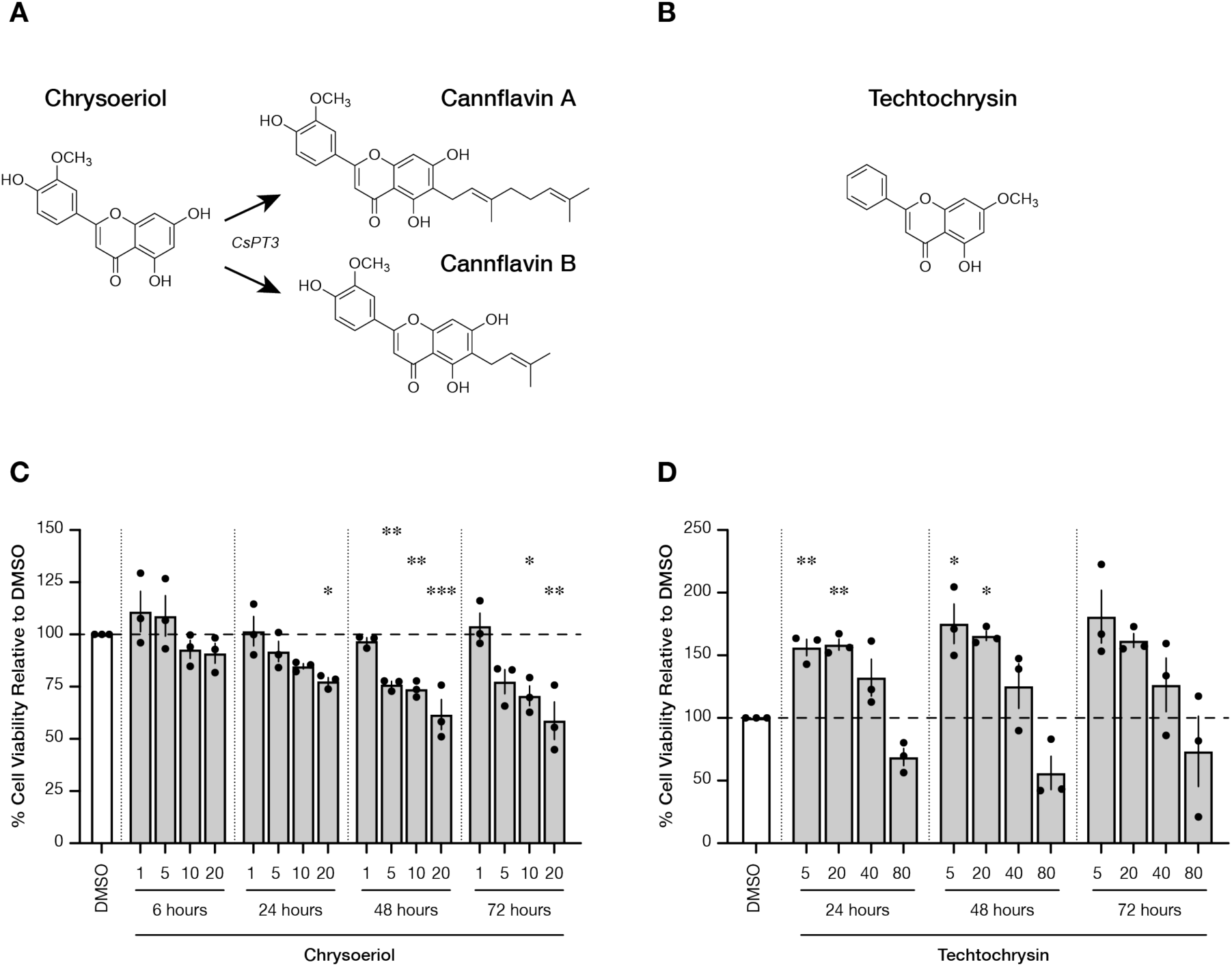
Effect of chrysoeriol and techtochrysin on U-87 GBM cell survival. **(A)** Chemical structure of chrysoeriol and **(B)** techtochrysin. As proposed by Real and colleagues (2019), *C. sativa* prenyltransferase 3 (CsPT3) catalyzes the biosynthesis of cannflavins from chrysoeriol. **(C-D)** U-87 cells were treated with increased concentrations of chrysoeriol (1-20 µM) or techtochrysin (5-80 µM), and cell viability was measured at the indicated time-points using the resazurin reduction assay. DMSO was used as a vehicle control. Application of chrysoeriol, but not techtochrysin, reduced the viability of U-87 GBM cells in a dose- and time-dependent manner. A one-way ANOVA was used to analyze the data from each cell line, treatment, and time independently with DMSO as the reference control. Tukey’s honestly significant difference (HSD) post hoc test: * *p* < 0.05, ** *p* < 0.01, *** *p* < 0.001. Graphs represent mean ± SEM (*n* = 3).

**Supplementary Figure 3.**
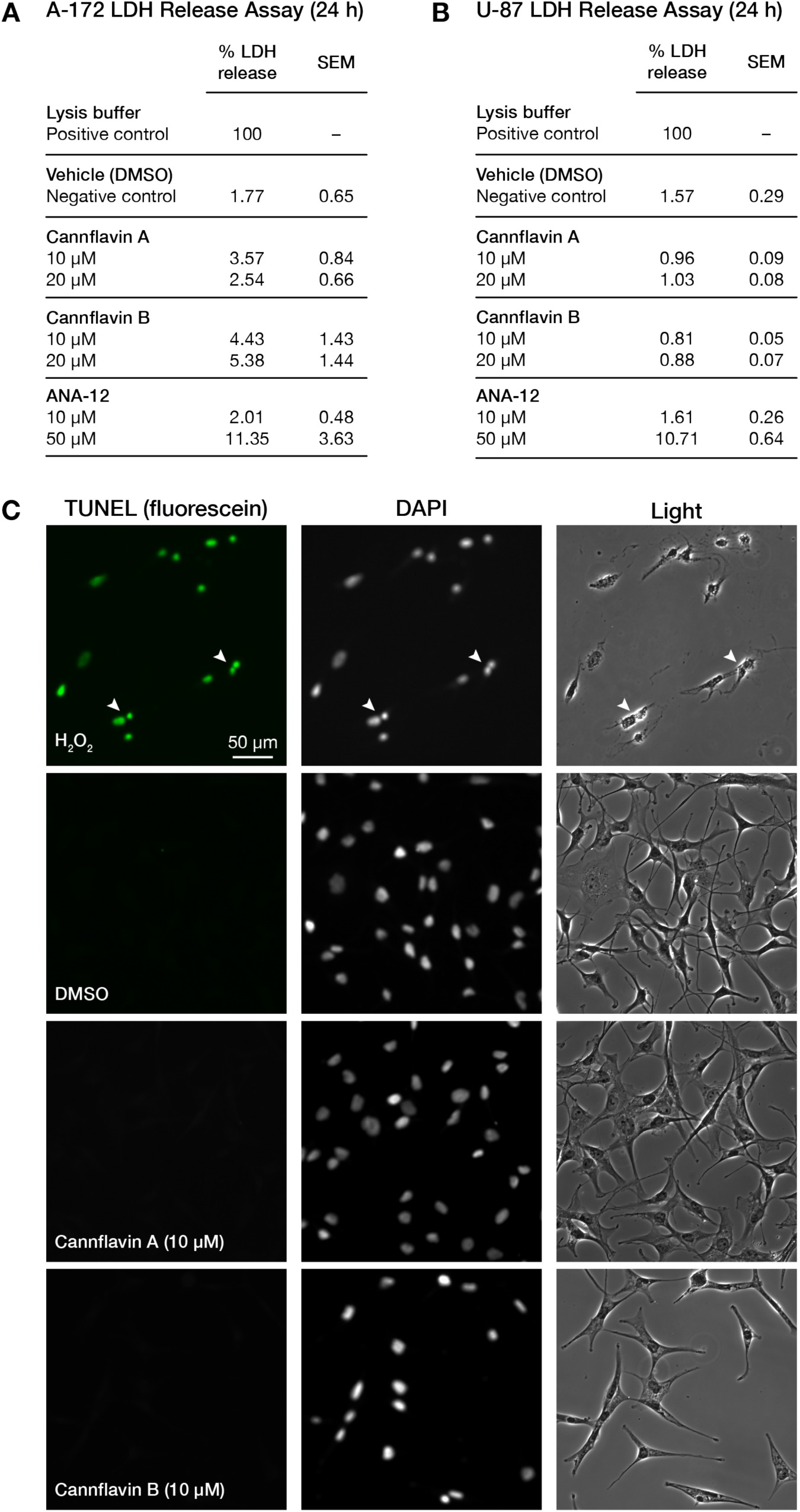
Effects of cannflavins A and B on LDH release and DNA fragmentation. **(A)** A-172 and **(B)** U-87 GBM cells were treated with cannflavin A or cannflavin B at 10 and 20 µM final concentrations, or ANA-12 (10 and 50 µM final concentration), and percent LDH release into cell culture media was quantified using the Invitrogen CyQUANT LDH Cytotoxicity Assay Kit. **(C)** Representative immunofluorescent images of A-172 GBM cells after 24 hours of treatment with cannflavins (10 µM final concentration) probed using TUNEL (fluorescein, green signal represent the occurrence of double-stranded DNA breaks). Nuclei were stained with DAPI (middle panels). Hydrogen peroxide (H_2_O_2_) was used as a positive control (applied directly to cell culture media for 10 minutes at a final concentration of 100 µM), and DMSO as the vehicle control for cannflavins. Arrowheads in the positive control condition show fragmented nuclei indicative of apoptosis. Scale bar = 50 µm.

**Supplementary Figure 4.**
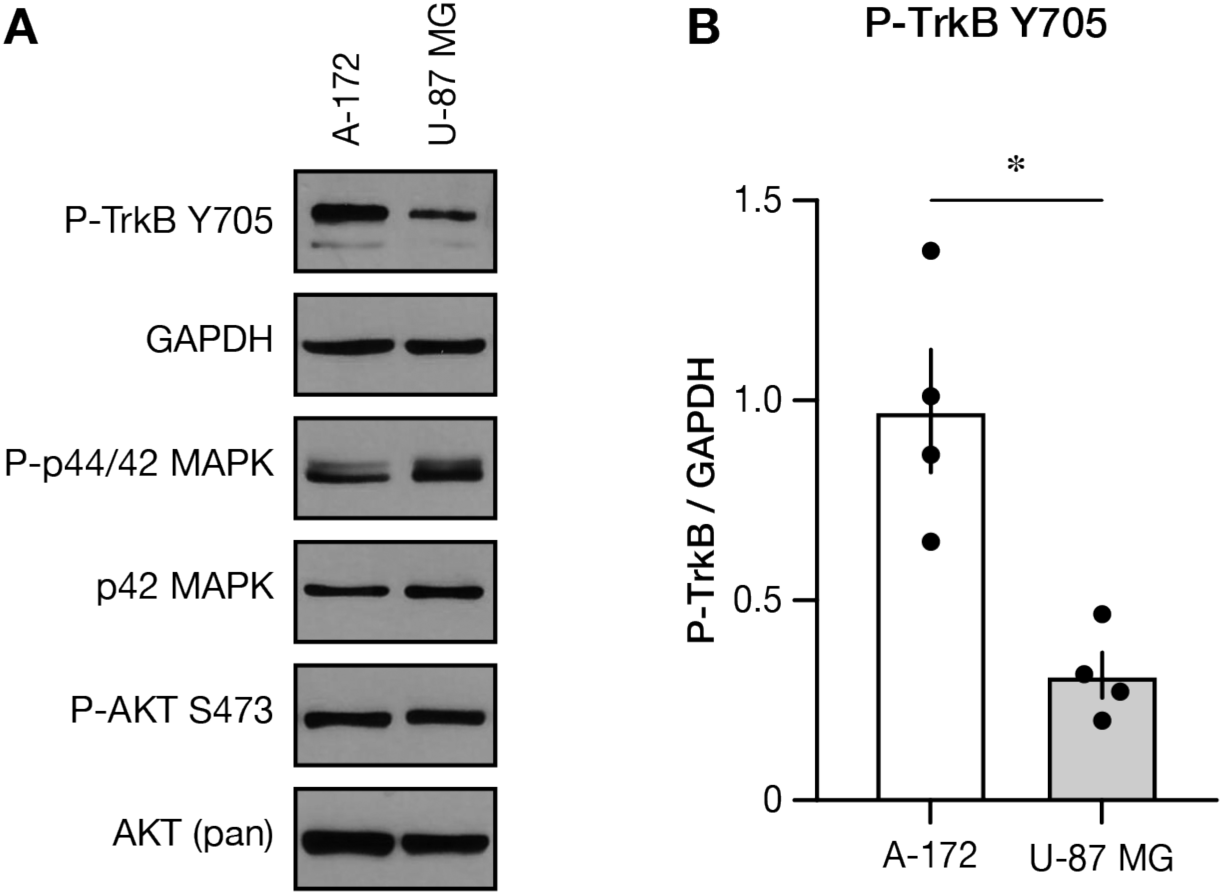
Level of phospho-TrkB Y705 between A-172 and U-87 untreated cells. **(A)** Representative western blots for phospho-TrkB Y705 and downstream signaling effectors. **(B)** Densitometry quantification shows a significantly higher abundance of phospho-TrkB Y705 in A-172 cells than in U-87 cells. * *p* < 0.05, two-tailed unpaired t-test. Graph represents mean ± SEM (*n* = 4).

**Supplementary Figure 5.**
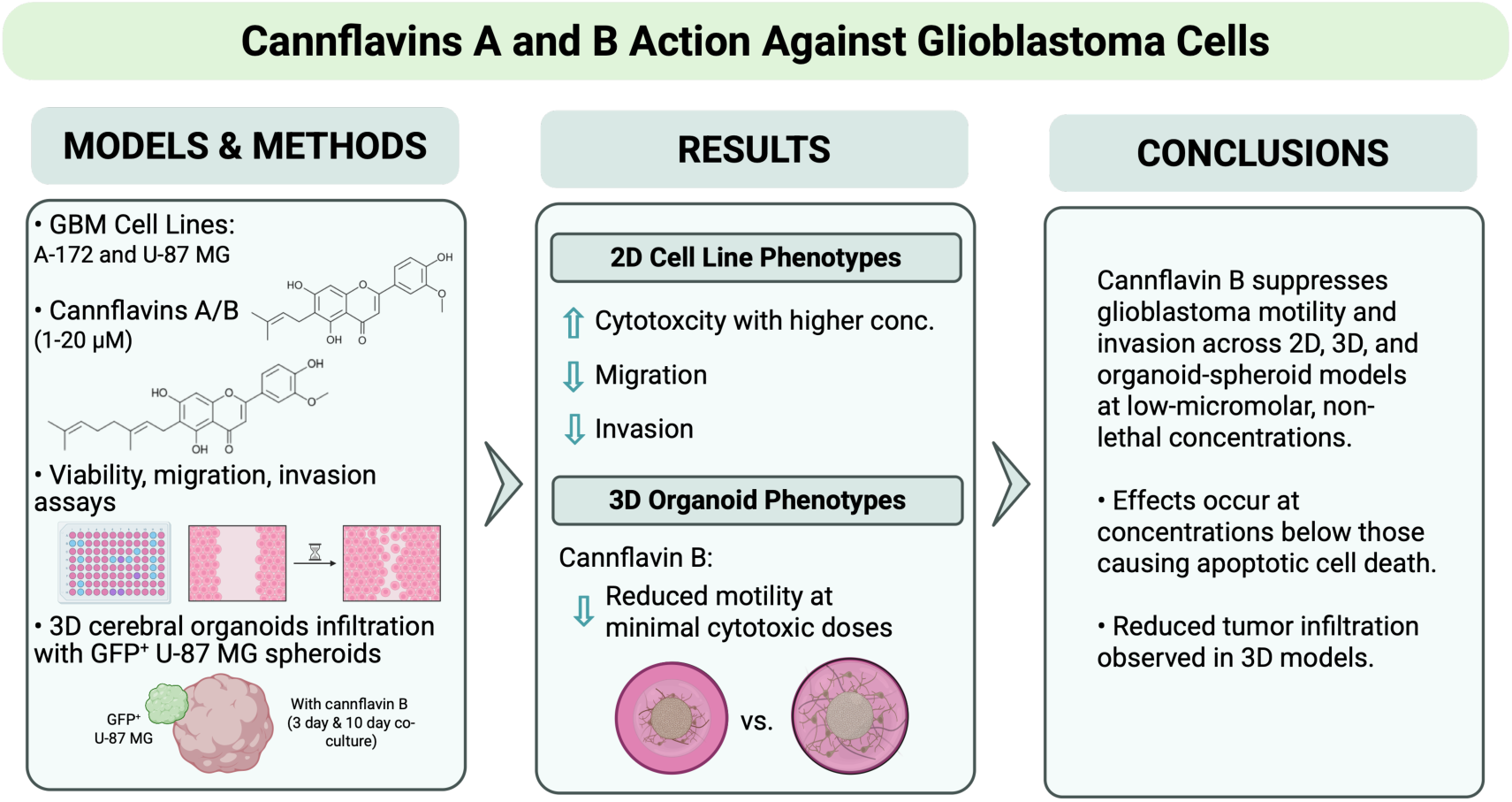
Graphical abstract.

**Supplementary Table.**
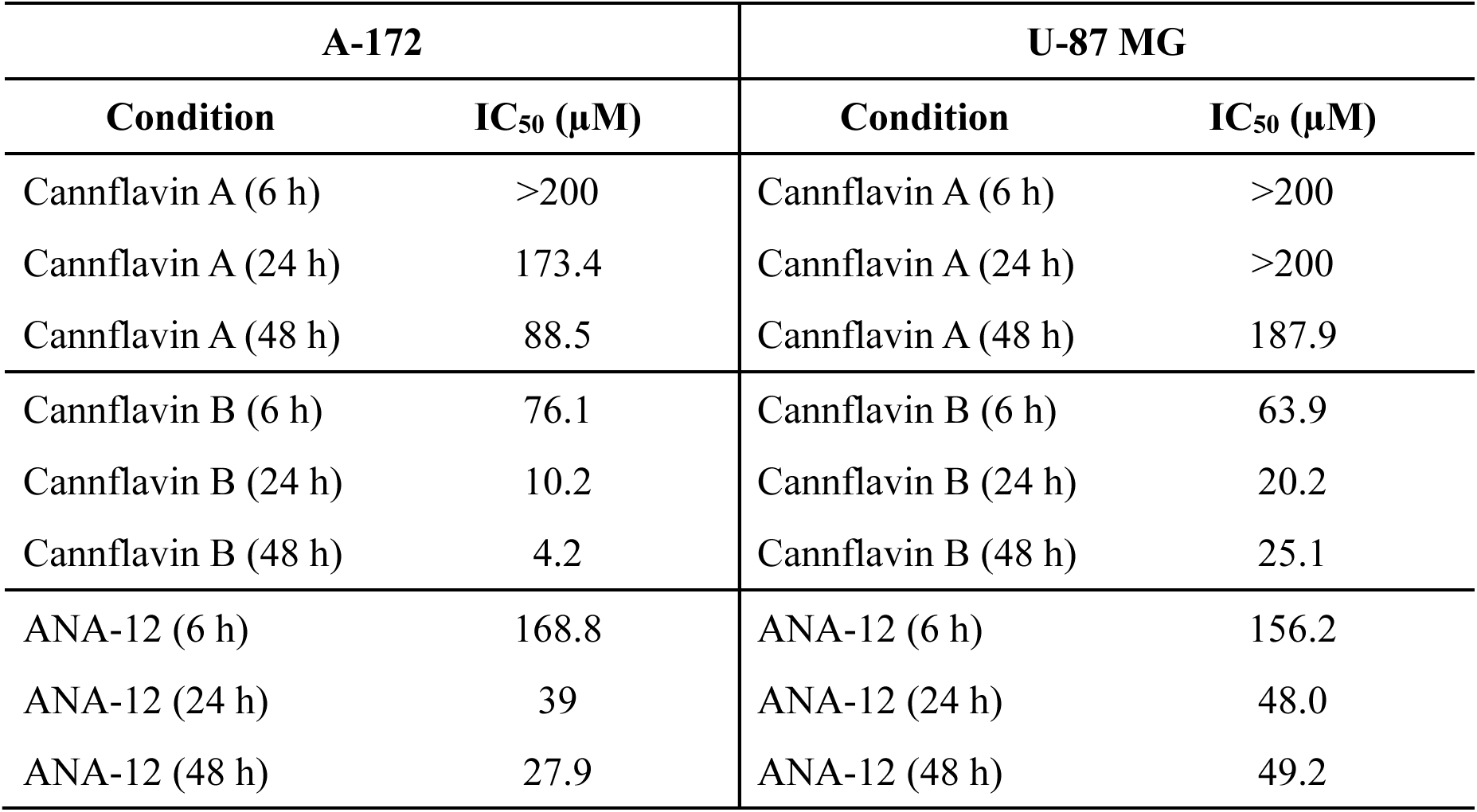
The estimated IC_50_ for the survival assay is shown in Figure 1.

**Supplementary Video.**
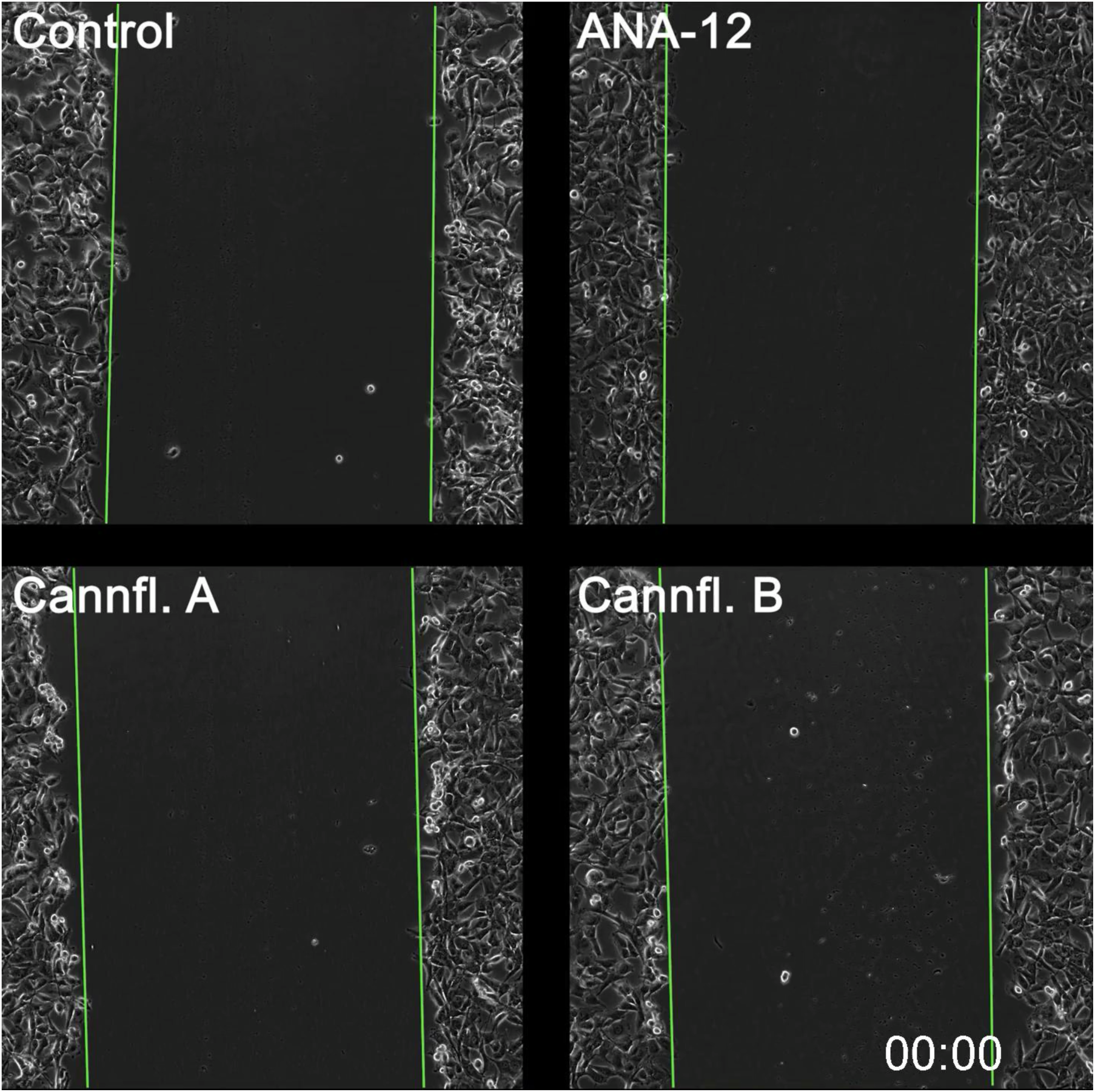
Scratch migration assay video. The green lines represent the initial boundaries (*t* = 0) of the scratched wound.

